# *Treponema denticola* dentilisin triggered TLR2/MyD88 activation upregulates a tissue destructive program involving MMPs via Sp1 in human oral cells

**DOI:** 10.1101/2021.01.18.427101

**Authors:** Sean Ganther, Allan Radaic, Nick Chang, Christian Tafolla, Ling Zhan, J. Christopher Fenno, Yvonne L. Kapila

## Abstract

Periodontal disease is driven by dysbiosis of the oral microbiome, resulting in over-representation of species that induce the release of pro-inflammatory cytokines, chemokines, and tissue-remodeling matrix metalloproteinases (MMPs) in the periodontium. These chronic tissue-destructive inflammatory responses result in gradual loss of tooth-supporting alveolar bone. The oral spirochete *Treponema denticola*, is consistently found at significantly elevated levels in periodontal lesions. Host-expressed Toll-Like Receptor 2 (TLR2) senses a variety of bacterial ligands, including acylated lipopolysaccharides and lipoproteins. *T. denticola* dentilisin, a surface-expressed protease complex comprised of three lipoproteins has been implicated as a virulence factor in periodontal disease, primarily due to its proteolytic activity. While the role of acylated bacterial components in induction of inflammation is well-studied, little attention has been given to the potential role of the acylated nature of dentilisin. The purpose of this study was to test the hypothesis that *T. denticola* dentilisin activates a TLR2-dependent mechanism, leading to upregulation of tissue-destructive genes in periodontal tissue. RNA-sequencing of periodontal ligament cells challenged with *T. denticola* bacteria revealed a significant upregulation of genes associated with extracellular matrix organization and degradation, including tissue-specific inducible MMPs that may play novel roles in modulating host immune responses yet to be characterized within the context of oral disease. The Gram-negative oral commensal, *Veillonella parvula*, failed to upregulate these same MMPs. Dentilisin-induced upregulation of MMPs was mediated via TLR2 and MyD88 activation, since knockdown of either TLR2 or MyD88 abrogated these effects. Challenge with purified dentilisin upregulated the same MMPs, whereas a dentilisin-deficient *T. denticola* mutant had no effect. Finally, *T. denticola*-mediated activation of TLR2/MyD88 led to the nuclear translocation of the transcription factor Sp1, which was shown to be a critical regulator of all *T. denticola-*dependent MMP expression. Taken together, these data support that *T. denticola* dentilisin stimulates tissue-destructive cellular processes in a TLR2/MyD88/Sp1-dependent fashion.

**AUTHOR SUMMARY:** Periodontal disease is driven by dysbiosis of the oral microbiome, which interacts with host tissues and thereby induces the release of pro-inflammatory cytokines, chemokines, and tissue-remodeling matrix metalloproteinases (MMPs), leading to destruction of the periodontal tissues. Even after clinical intervention, patients with severe periodontal disease are left with a persistent pro-inflammatory transcriptional profile throughout the periodontium. The oral spirochete, *Treponema denticola*, is consistently found at elevated levels in periodontal lesions and is associated with several pathophysiological effects driving periodontal disease progression. The *T. denticola* surface-expressed protease complex (dentilisin) has cytopathic effects consistent with periodontal disease pathogenesis. To date, few direct links have been reported between dentilisin and the cellular and tissue processes that drive periodontal tissue destruction at the transcriptional and/or epigenetic levels. Here, we utilize wild type and dentilisin-deficient *T. denticola* as well as purified dentilisin to characterize dentilisin-dependent activation of intracellular pathways controlling MMP expression and activity. Our results define a role for dentilisin in initiating this signal cascade. Also, our study identified tissue-specific inducible MMPs that may play novel roles in modulating as-yet uncharacterized host responses in periodontal disease. Lastly, *T. denticola* dentilisin stimulates tissue-destructive cellular processes in a TLR2/MyD88/Sp1-dependent fashion. Taken together, our study provides new insights into the molecular mechanisms underpinning periodontal disease progression which could lead to the development of more efficacious therapeutic treatments.

## INTRODUCTION

Periodontitis or Periodontal Disease is characterized as a chronic oral inflammatory disease that compromises the integrity of the tooth-supporting tissues, which include the gingiva, periodontal ligament and alveolar bone, and are collectively known as the periodontium. In its severe form, which afflicts 8.5% of adults in the United States(1), periodontitis may cause tooth loss, and also affect systemic health by increasing the patients’ risk for atherosclerosis, adverse pregnancy outcomes, rheumatoid arthritis, aspiration pneumonia and cancer(2-5). Additionally, patients with chronic forms of periodontal disease are left with a non-resolving pro-inflammatory transcriptional profile throughout the periodontium, even after clinical intervention, leading to tissue-destruction and tooth loss(6-8). This suggest that previously uncharacterized cellular and molecular mechanisms underlying periodontal disease pathophysiology may explain why many patients do not respond to the conventional treatment schema.

The periodontal ligament (PDL) has two primary functions: 1) to absorb and respond to mechanical stresses, and 2) to provide vascular supply and nutrients to the cementum, alveolar bone and the PDL itself. Osteoblasts, osteoclasts, cementoblasts and endothelial cells make up the PDL and reside on the surface of the lamina dura and endosteal surfaces of the alveolar bone and cementum(9, 10). The most prominent cell type are human periodontal ligament fibroblasts (hPDL), which specialize in mechanosensing and tissue remodeling through robust expression of various hydrolytic enzymes such as matrix metalloproteinases (MMPs)(10, 11). They also function as immune-like cells demarcated by the production of inflammatory cytokines, chemokines and expression of pattern recognition receptors (PRRs), such as toll-like receptors (TLRs), which are responsible for monitoring the local environment for signs of danger in the form of highly conserved microbial molecules present during periodontal disease (12-15). When TLR signaling is left unchecked, downstream genes, such as MMPs can become significantly upregulated and constitutively activated, thereby contributing to the overall destruction of the periodontium by degrading extracellular matrix (ECM) proteins, such as collagen and fibronectin(11, 16-20). Although the role of MMPs has been primarily ascribed to turnover of the ECM, the great number of new substrates discovered for MMPs in the last few years suggest they are capable of regulating many signaling pathways, cell behaviors and diseases through novel mechanisms(21). Therefore, PDL cells likely play a significant role in initiating and exacerbating tissue degradation and suggest that pharmacological anti-inflammatory treatment of periodontal disease should target dysregulated MMPs activity and levels in the PDL tissues.

The pathogenic processes of periodontal disease are primarily due to the host response, which propagates the destructive responses initiated by microbes(22-24). A triad of oral anaerobic bacteria called the “Red Complex” comprised of *Porphorymonas gingivalis, Treponema denticola* and *Tannerella forsythia*, have traditionally been considered as causative agents of periodontitis, based on their virulence properties and strong association with diseased sites(25-27). While a large focus has been placed on species such as *P. gingivalis, Fusbacterium nucleatum, and Aggrigatibacter actinomycetemcomitans*, recent metagenomic and metatranscriptomic studies have implicated *T. denticola* in advanced or aggressive forms of periodontal disease and recurrence of disease(25, 28-42). Greatly increased numbers of *T. denticola* levels in periodontal disease vs health has been well-documented in the literature for >40 years(43). Our lab has readily demonstrated that *T. denticola* was ∼15-fold higher in the subgingival biofilm of periodontal lesions(44). Additionally, elevated *T. denticola* biofilm levels combined with elevated MMP levels in host tissues display robust combinatorial characteristics in predicting advanced periodontal disease severity(44, 45). Further, Lee *et al*. 2009 and colleagues demonstrated that *T. denticola* infection was sufficient to induce alveolar bone resorption in a mouse model of periodontal disease(46). Thus, clinical data regarding the increased presence of *T. denticola* in periodontal lesions, together with basic and *in vivo* studies involving the role of *T. denticola* products suggest that it plays a pivotal role in driving periodontal disease progression.

Among the various *T. denticola* effector molecules that have been described, its acylated chymotrypsin-like protease complex (CTLP), more recently called dentilisin, is a major virulence factor which facilitates numerous cytopathic effects that align with periodontal disease pathophysiology(32, 47, 48). A few examples including adhesion, degradation of endogenous ECM-substrates(37), tissue penetration(49), complement evasion(50), ectopic activation of host MMPs(41) and degradation of host chemokines and cytokines, such as IL-1*β* and IL-6, primarily due to its potent proteolytic activity(40). However, as in other spirochetes, conserved lipid moieties of the protease complex recognized by host TLR2 receptor complexes may contribute to activation of innate immune responses(51). Because predominant host responses to lipoproteins are believed to be to their lipid moieties, most studies have focused on diacylated lipopeptide, Pam2CSK4, and triacylated lipopeptide, Pam3CSK4, which mimic bacterial lipoproteins for their potent immunostimulatory and osteoclastogenic activities by preferentially activating TLR2-dependent pathways(51-53). Recent studies have demonstrated that synthetic di- and tri-acylated lipopeptides which preferentially activate TLR2/6 and TLR2/1-dependent pathways respectively, are sufficient to induce alveolar bone loss in mice(52, 54), broadening the avenues of investigation into the role of lipoproteins underpinning the pathogenesis of periodontal disease. However, studies which utilize endogenously expressed bacterial lipopeptides are greatly lacking.

While a clear role for dentilisin in the context of periodontal disease has been delineated at the protein level, its ability to influence host ECM metabolism in a gene-centric context has yet to be characterized. Thus, the aim of this study was to determine the extent to which *T. denticola* and its highly expressed acylated dentilisin protease complex influence the transcriptional regulation of MMPs through TLR2-dependent pathways in hPDL cells.

## RESULTS

### *T. denticola* upregulates genes associated with ECM and degradation in hPDL cells

To better understand the extent to which *T. denticola* influences host transcriptomes in an unbiased and gene-centric context, we challenged three healthy patient replicates of hPDL cells with wild-type Td-WT for 2 hours. The cells were then incubated for an additional 3 or 22 hours (5 and 24-hours, respectively) before extracting total RNA for sequencing. Unchallenged hPDL cells were used as a negative control. The results are shown in **Figure 1**. In order to include moderately up- or downregulated genes in the analysis, genes below our cutoff criterion with a significant difference (*p* < 0.05) were examined. Differential expression analysis revealed a large overlap in gene expression between the control and 5-hour incubation groups (289 genes), whereas the control and 24-hour incubation groups showed significantly less overlap (165 genes) **(Figure 1A)**. Hierarchical clustering analysis was carried out with the log10 (FPKM+1) of union differential expression genes of all comparison groups under different experimental conditions. Essentially, experimental groups which have more unique clusters diverge further away from the control group. The 24-hour incubation group clustered away from the 5-hour incubation and control groups **(Figure 1B)**. For the gene ontology analysis, we utilized the mean FPKM values for each gene across experimental groups. Gene clusters at or above a 0.05 adjusted p-value were delineated as statically significant increases. ECM (∼6-fold change), collagen degradation (∼2-fold change) and degradation of the ECM (∼2-fold change) amongst the top 20 significantly enriched biological processes **(Figure 1C)**.

**Figure 1.**
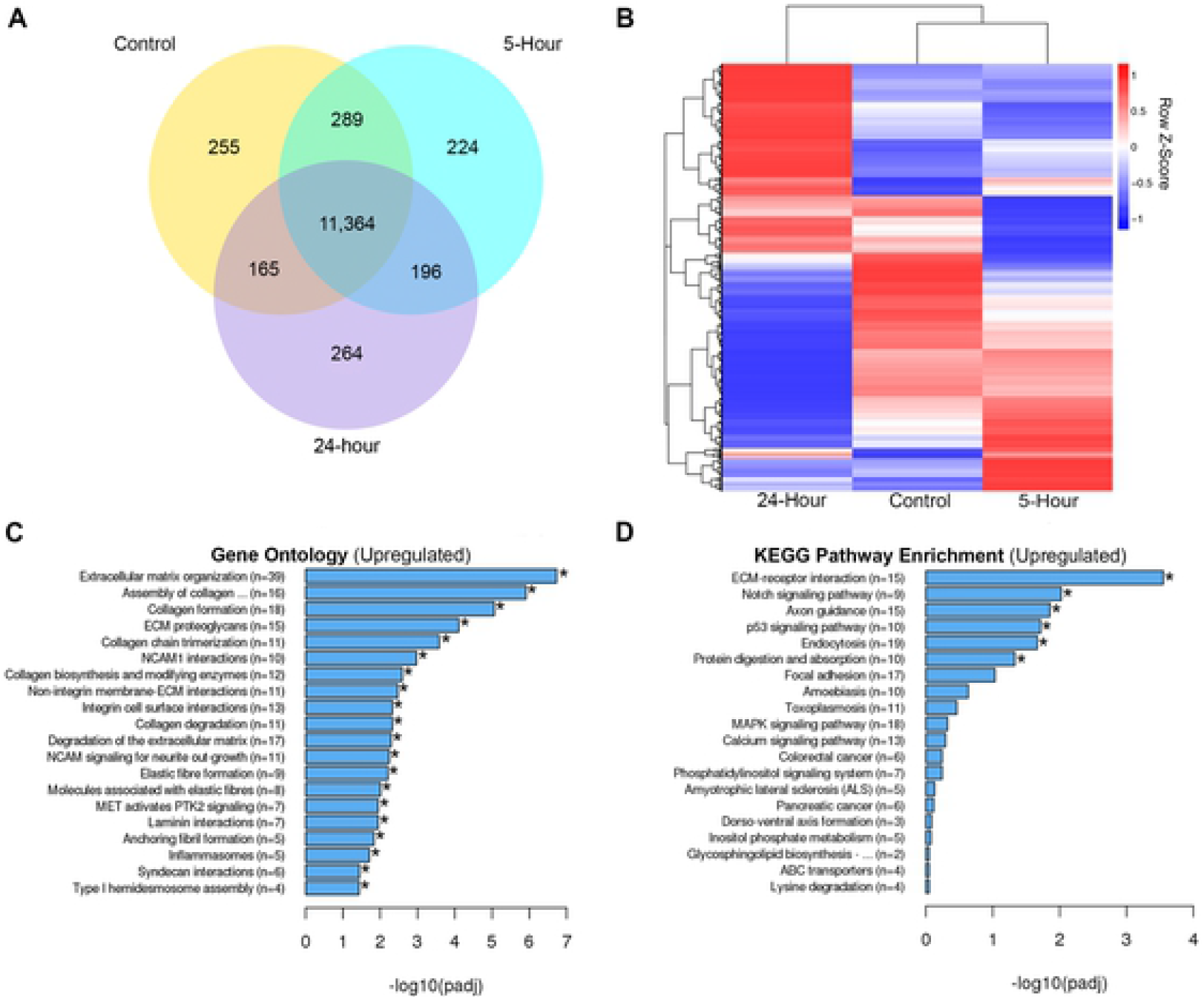
Differential expression analysis of *T. denticola* challenged hPDL cells. Total RNA was extracted from healthy patient-derived hPDL cells challenged with Td-WT bacteria at a MOI of 50 for 2-hours in media free of supplements followed by 3 and 22-hours in media supplemented with 10% FBS, 1% Pen Strep and 1% Amphotericin B. Mean FPKM values were used for downstream analysis (n=3 patient replicates). A) Overlapping and differentially expressed genes between control, 5-hour and 24-hour incubation groups visualized using a Venn diagram. B) Hierarchical clustering analysis was used to determine similarity of transcriptome profiles based on differential expression as a heatmap. Red (Upregulation) to blue (Downregulation) color gradient of heatmap represents normalized gene expression as row Z-scores. C) Top 20 enriched Gene Ontology terms of hPDL cells challenged for 2-hours followed by a 22-hour incubation using the Reactome nomenclature. Statistical significance was assessed using a Kolmogorov-Smirnov test followed by Benjamini-Hochberg correction (p<0.05). D) Top 20 enriched signaling pathways of hPDL cells challenged for 2-hours followed by a 22-hour incubation using the Kyoto Encyclopedia of Genes and Genomes (KEGG) database. Statistical significance was assessed using a Kolmogorov-Smirnov test followed Benjamini-Hochberg correction (p<0.05).

Downregulated GO and KEGG Terms can be found in **Supplemental Figure 1**. KEGG pathway analysis revealed ECM-Receptor (∼4-fold change), Notch (∼2-fold change), endocytosis (∼2-fold change) and protein digestion (∼2-fold change) signaling amongst the top 20 enriched signaling pathways upon *T. denticola* challenge **(Figure 1D)**. Interestingly, Membrane-type matrix metalloproteinase 4 (MMP-17), Stromelysin-3 (MMP-11) and Epilysin (MMP-28) were consistently among the most highly expressed genes clustered with various GO terms associated with ECM remodeling and catabolism (Data not shown) and have not been associated with periodontal disease to date. Although RNA-seq analysis revealed increases in MMP-2 and MMP-14 read counts, they were not found to be statistically significant (data not shown). Both genes, however, have been found to be chronically upregulated in hPDL fibroblasts upon *T. denticola* challenge for up to 12 days *in vitro*(44). Additionally, while the primary functions of these enzymes has been associated with tissue turnover, both MMP-2 and MMP-14 participate in the active degradation of various cytokines(55), chemokines(55), ECM proteins(56) and various intracellular substrates(21, 57, 58) expressed by hPDL cells expressed during periodontal disease infections, delineating their potentially important role in driving periodontal induced tissue destruction. Next, we sought to determine if this upregulation of tissue destructive genes is common among pathogens and commensal bacteria.

### MMPs are not upregulated by commensal oral Gram-negative Veillonella parvula

To determine if the induction of these MMPs was specific to *T. denticola*, we utilized *Veillonella parvula*, a Gram-negative anaerobic oral commensal that is prevalent in the gut and oral cavity and is not associated with oral diseases(59). hPDL cells were challenged with wild-type *T. denticola* or *V. parvula* bacteria for 2 hours at 50 MOI followed by a 22-hour incubation period in *α*-MEM media free of antibiotics and supplemented with 10% FBS. *V. parvula* challenge did not induce the upregulation of MMPs *2, 11, 14, 17* or *28* **(Figure 2)**. By contrast, *T. denticola* elicited a ∼2-fold upregulation or more of all target MMP genes **(Figure 2)**.

**Figure 2.**
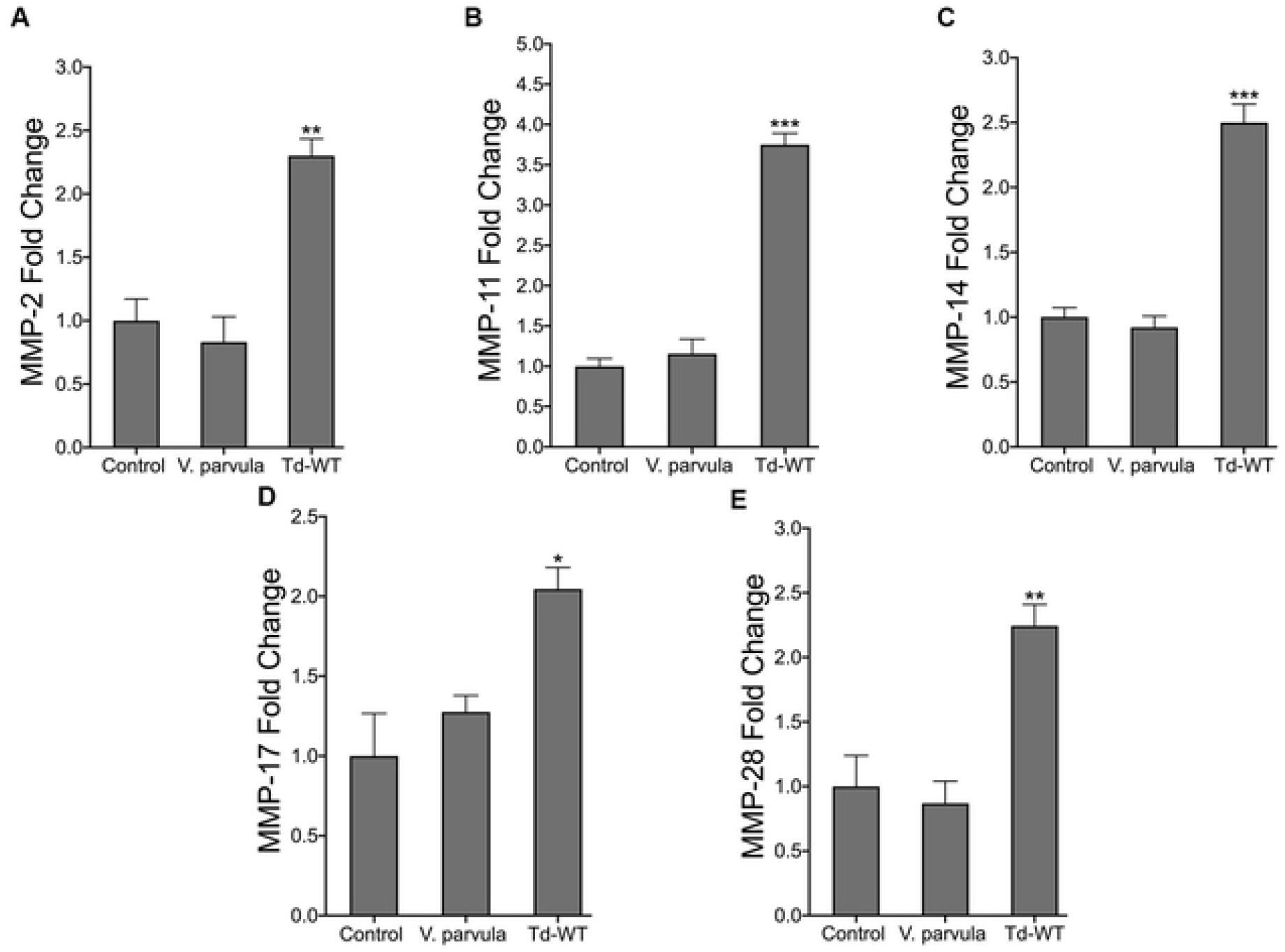
*T. denticola* upregulates MMPs in hPDL cells while *V. parvula* does not. (A-E) RT-qPCR for *MMP-2, MMP-11, MMP-14, MMP-17 and MMP-28* mRNA expression of healthy hPDL cells challenged with *V. parvula (*ATCC 10790) and Td-WT bacteria at a MOI of 50 for 2-hours, followed by a 22-hour incubation in media supplemented with 10% FBS and 1% Pen/Strep and 1% Amphotericin B. Expression of each gene was normalized to that of *GAPDH*. Statistical significance was determined using a One-Way ANOVA followed by Post-Hoc Tukey’s multiple comparisons. Bars represent mean ± SEM (n=5). ^*^*p*<.05 versus control. ^**^*p<*.01 versus control. ^***^*p<*.001 versus control.

### *T. denticola* dentilisin mediates the upregulation of various MMPs in hPDL cells

Among the various TLR2 stimulatory bacterial ligands, lipoproteins are considered as a major virulence factor because of their strong immunostimulating capacity(53, 60, 61). Interestingly, while differential innate immune responses have been reported depending on the acylation status and tissue type(61-63), both di- and triacylated synthetic lipopeptides have demonstrated the ability to sufficiently drive alveolar and long bone resorption in mice through TLR2/MyD88 dependent mechanisms(52, 54). We investigated whether purified dentilisin can drive MMP gene regulation in hPDL cells and the results can be seen in **Figure 3**. hPDL cells were challenged with dentilisin purified from *T. denticola* ATCC 35405 at a final concentration of 1 *µ*g/mL for 2-hours in MEM-media supplemented 1% P/S. Enzymatic activity of dentilisin protein was verified by collecting conditioned media from dentilisin stimulated hPDL cells as described above and analyzing samples using gelatin zymography **(Supplemental Figure 2)**. While, dentilisin enzymatic activity (98 kDa) increased in a dose dependent manner, MMP-2 enzymatic activity (72 kDa) concomitantly increased in a statistically significant manner (∼2.0-fold) across all 3 experimental conditions.

**Figure 3.**
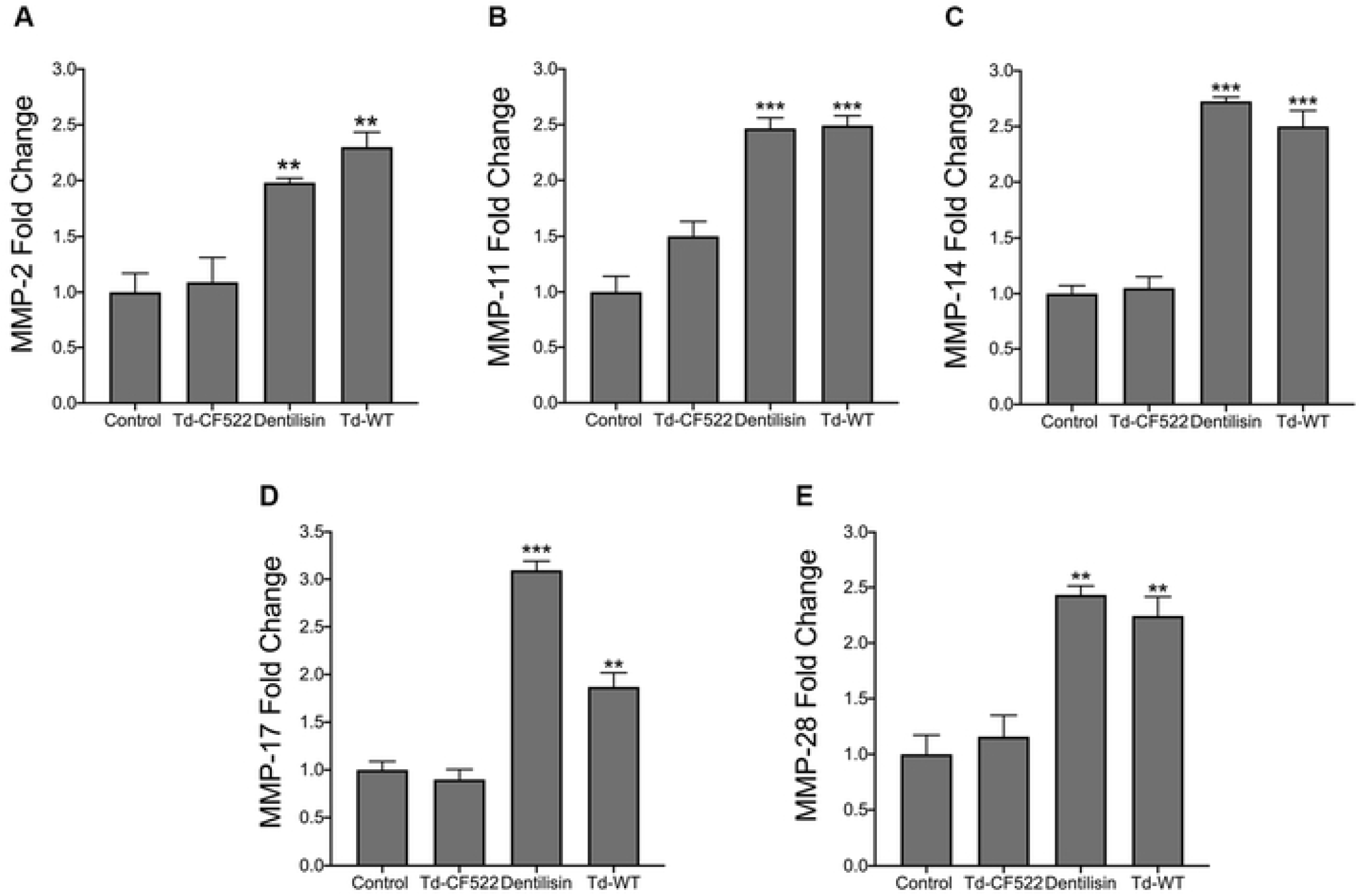
*T. denticola s*urface-expressed dentilisin mediates the upregulation of MMP 2, 11, 14, 17 and 28 mRNA levels in hPDL cells. (A-E) RT-qPCR for *MMP-2, MMP-11, MMP-14, MMP-17 and MMP-28* mRNA expression of healthy hPDL cells challenged with isogenic Td-CF522 bacteria, Td-WT bacteria and purified dentilisin. Cells were stimulated at an MOI of 50 and a final concentration of of 1 ug/mL. Cells were challenged for 2-hours in alpha-MEM media supplemented with 10% FBS followed by a 22-hour incubation in alpha-MEM media with 10% FBS, 1% PenStrep and 1% Amphotericin B. The expression of each gene was normalized to that of *GAPDH*. Statistical significance was determined using a One-Way ANOVA followed by Tukey’s Post-Hoc multiple comparisons. Bars represent mean ± SEM (n=4). ^**^*p<*.01 versus control. ^***^*p<*.001 versus control.

Total RNA was extracted and processed for cDNA synthesis from Td-WT, Td-CF522 or purified dentilisin challenged/stimulated hPDL cells and rendered to RT-qPCR. Cells stimulated with purified dentilisin resulted in statistically significant increases in all MMP targets with MMP-17 being the most responsive **(Figure 3)**. Similarly, hPDL cells challenged with Td-WT bacteria resulted in statistically significant increases across all MMP targets (∼2-2.5-fold) **(Figure 3)**. By contrast, hPDL cells challenged with dentilisin-deficient *T*.*denticola* (Td-CF522) resulted in no statistically significant upregulation across all MMP targets compared to the control group **(Figure 3)**.

### shRNA knockdown of *TLR2* inhibits *T. denticola*-induced upregulation of *MMPs 2, 14 17 and 28* in hPDL cells while synergistically increasing MMP 11 expression

Thus, after determining dentilisin plays an important role as a regulator of MMP expression in host hPDL cells, we sought to establish a direct link between TLR2 activation and MMP gene regulation. TLRs, particularly *TLR2*, and to a much lesser extent *TLR4*, regulate important immune responses to periodontal bacteria *in vivo* and *in vitro(36, 64-70)*. TLR2 has been identified as a receptor for an array of ligands comprising *T. denticola* bacteria, including lipoteichoic acid, beta barrel proteins, and acylated lipopeptides, such as dentilisin(71, 72). Thus, to examine the role of TLR2 and MMP transcriptional regulation in hPDL cells, we generated a primary hPDL *TLR2* knockdown cell line using shRNA targeted against the TLR2 gene and transduced using lentiviral particles. Basal *TLR2* expression was reduced ∼60% across 3 clonal replicates as validated using RT-qPCR; reduction was found to be statistically significant using an unpaired t-test **(Figure 4A)**. Next, *TLR2* knockdown cells were challenged with either purified dentilisin or wild-type *T. denticola* bacteria. *MMP 2, 11, 14, 17 and 28* expression increased upon stimulation with both purified dentilisin and Td-WT in the shRNA control group **(Figure 4B-4F)**. Compared with hPDL cells transfected with the scrambled shRNA control, induction of expression of MMPs 2, 14, 17 and 28 was suppressed in *TLR2*-deficient hPDL cells challenged with *T. denticola* or dentilisin, while MMP 11 was significantly upregulated in *T. denticola*-challenged *TLR2*-deficient hPDL cells. After determining TLR2 activation plays an integral role in the regulation of MMP expression, we next sought out to determine which downstream mediator is required to propagate this signal.

**Figure 4.**
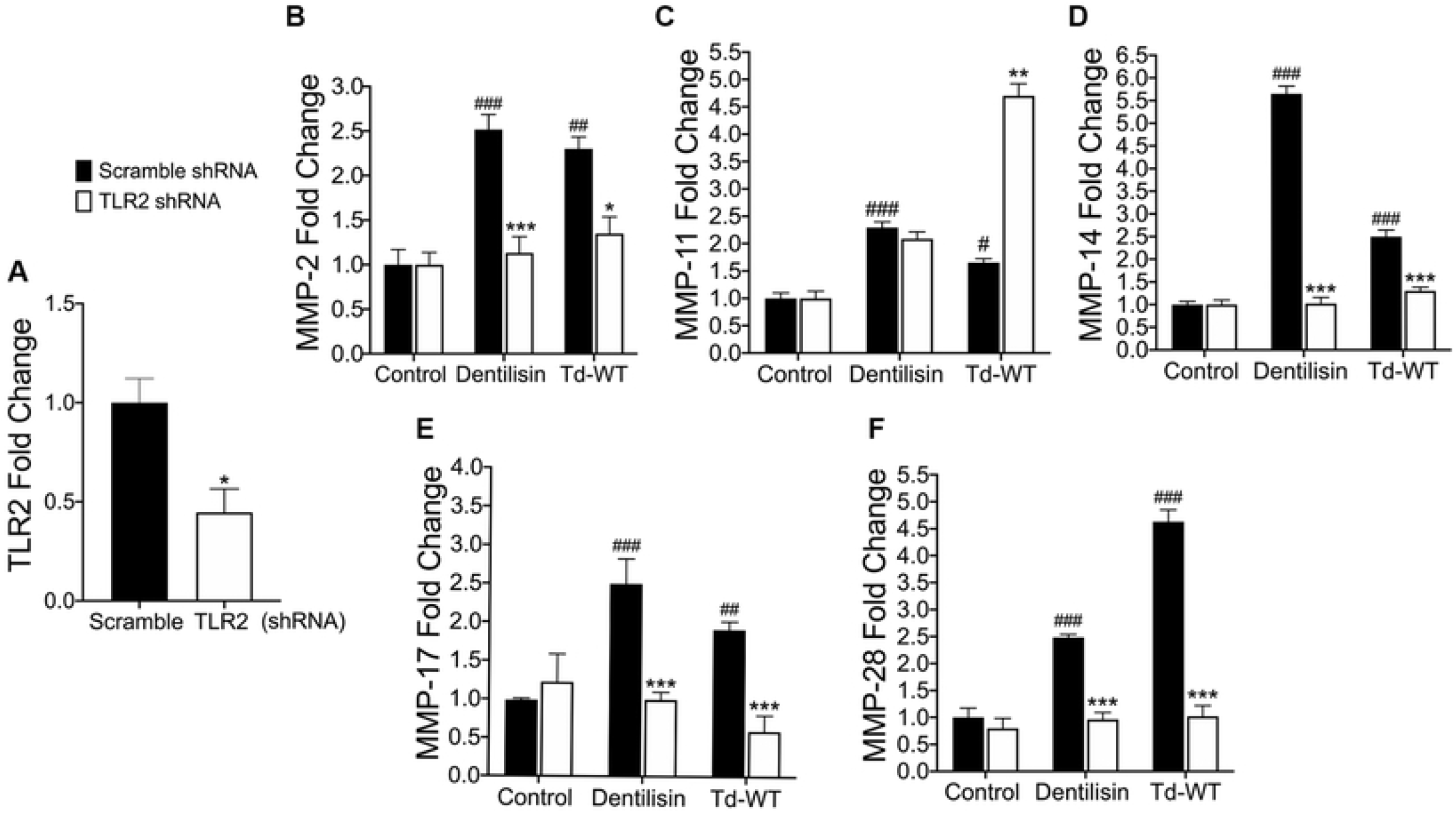
Suppression of *TLR2* inhibits *T. denticola*-stimulated upregulation *of MMPs 2, 14, 17* and *28* while exacerbating MMP *11* expression in hPDL cells. A) RT-qPCR validation of stable gene suppression using shRNA vectors targeted against *TLR2* in healthy hPDL cells. Cells transduced with scramble shRNA vectors were used as a control. Statistical significance was determined using an unpaired t-test. Bars represent ± SEM of mean value (n=3 clones). ^*^*p*<.05 versus control. B-F) RT-qPCR for *MMP-2, MMP-11, MMP-14, MMP-17 and MMP-28* mRNA expression of scramble shRNA control and *TLR2* shRNA hPDL cells challenged or stimulated with Td-CF522, purified dentilisin or Td-WT at an MOI of 50 and concentration of 1 *µ*g/mL for 2-hours in alpha-MEM media with no supplementation followed by a 22-hour incubation in alpha-MEM media supplemented with 10% FBS, 1% Pen/Strep and 1% Amphotericin B. The expression of each gene was normalized to that of *GAPDH*. Statistical significance was determined using a Two-Way ANOVA followed by post-hoc Tukey’s multiple comparisons. Bars represent mean ± SEM (n=3). #*p*<.05 versus scramble control group. ##*p*<.01 versus scramble control group. ###*p*<.001 versus scramble control group. ^*^*p<*.05 versus *TLR2* shRNA equivalent group. ^***^*p<*.001 versus *TLR2* equivalent shRNA group.

### shRNA knockdown of *MyD88* inhibits *T. denticola*-stimulated upregulation of *MMPs 2, 11, 14, 17 and 28* in hPDL cells

All TLRs signal through MyD88 with the exception of TLR3 and TLR4 which utilize TIRAP(73). However, numerous recent reports suggest that MyD88-independent TLR2 pathways may also contribute to periodontal disease progression(74). Thus, to determine if *T. denticola*-induced TLR2 activation requires MyD88 to regulate targeted MMPs, we generated an hPDL *MyD88* knockdown cell line, achieving approximately ∼60% reduction in *MyD88* basal expression across 3 validated clonal replicates **(Figure 5A)**. *MyD88*-deficient hPDL cells were stimulated with either Td-WT or purified dentilisin as described above. In the negative shRNA control group, all MMP targets showed increased mRNA levels following challenge with both Td-WT and dentilisin **(Figure 5B-5F)**. By contrast, *MyD88*-deficient cells challenged with Td-WT bacteria and purified dentilisin revealed unanimous depression of all MMP targets as compared to the respective scramble shRNA control samples **(Figure 5B-5F)**.

**Figure 5.**
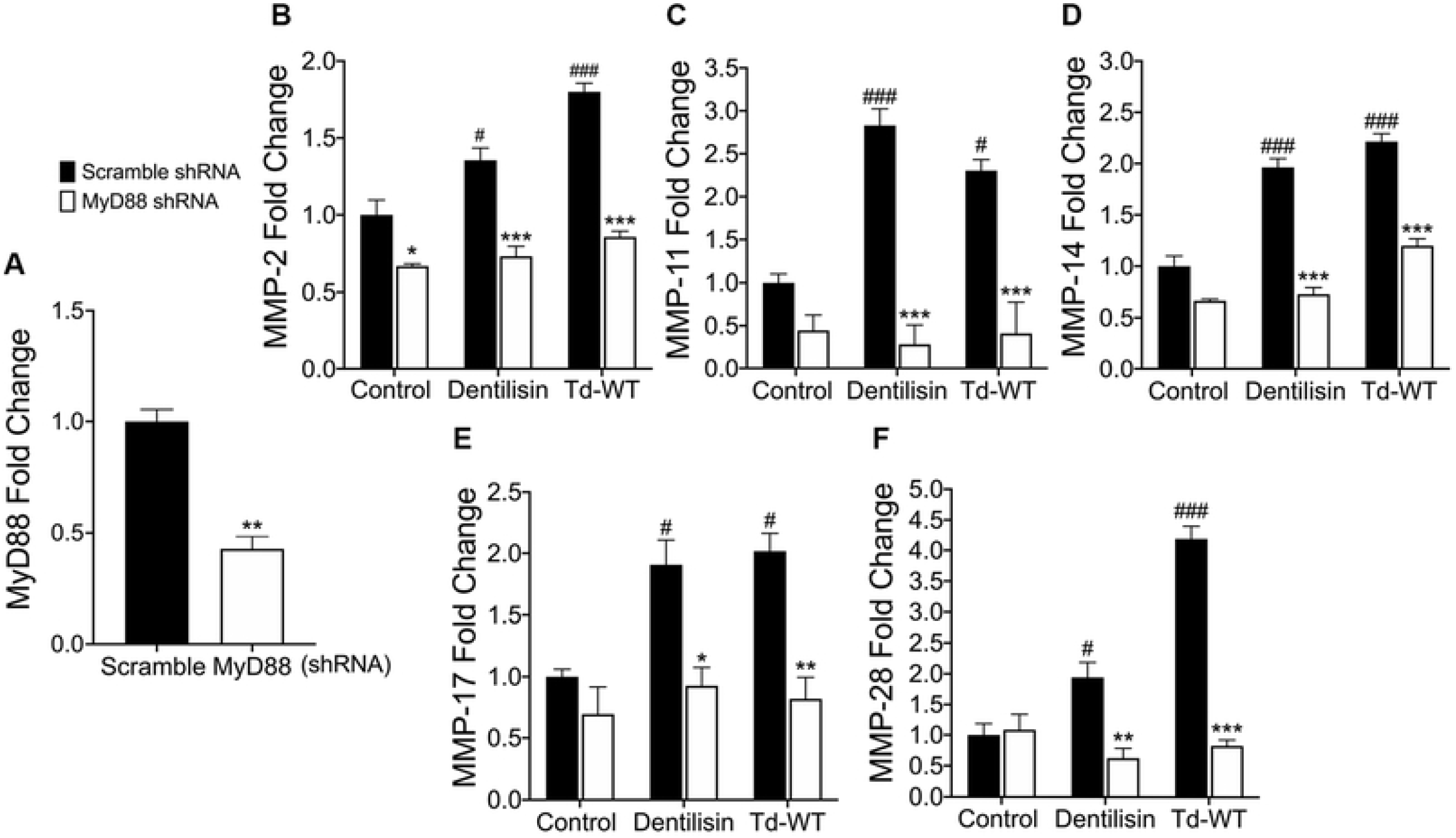
Suppression of *MyD88* inhibits *T. denticola*-stimulated upregulation of MMP *2, 11, 14, 17* and *28* in hPDL cells. A) RT-qPCR validation of stable gene suppression using shRNA vectors targeted against *MyD88* in healthy hPDL cells. Cells transduced with scramble shRNA vectors were used as a control. Statistical significance was determined using an unpaired t-test. Bars represent ± SEM of mean values (n=3 clones). ^*^*p*<.05 versus control. B-F) RT-qPCR for *MMP-2, MMP-11, MMP-14, MMP-17 and MMP-28* mRNA expression of scramble shRNA control and *MyD88* shRNA hPDL cells challenged or stimulated with Td-CF522, purified dentilisin or Td-WT at an MOI of 50 and concentration of 1 *µ*g/mL for 2-hours in alpha-MEM media with no supplementation followed by a 22-hour incubation in alpha-MEM media supplemented with 10% FBS, 1% Pen/Strep and 1% Amphotericin B. The expression of each gene was normalized to that of *GAPDH*. Statistical significance was determined using a Two-Way ANOVA followed by post-hoc Tukey’s multiple comparisons. Bars represent mean ± SEM (n=3). #*p*<.05 versus scramble control group. ###*p<*.001 versus scramble control group ^*^*p*<.05 versus *MyD88* shRNA equivalent group. ^**^*p<*.01 versus *MyD88* shRNA equivalent group. ^***^*p<*.001 versus *MyD88* shRNA equivalent group.

### shRNA knockdown of TLR2/MyD88-dependent signaling inhibits nuclear translocation of transcription factor Sp1 in hPDL cells stimulated with *T. denticola* or *dentilisin*

One of the many downstream targets of TLR/MyD88 signaling is the transcription factor Specificity Protein 1 (Sp1). Sp1 regulates a variety of genes associated with cell cycle, pro-inflammation and tissue-destruction, including *MMPs 2, 11, 14 17 and 28*. Interestingly, Larsson and colleagues have readily demonstrated that mRNA levels of Sp1 are consistently upregulated at periodontally involved sites in humans(75). Thus, to determine if *T. denticola* influences Sp1 expression in the PDL, hPDL cells were challenged with Td-WT and Td-CF522 bacteria at a multiplicity of infection of 50 for 2-hours followed by a 22-hour incubation period in media. Cells were collected and used for western blot analysis utilizing a monoclonal antibody against total Sp1 protein. Total Sp1 protein expression levels increased in the groups challenged with Td-WT at 50 MOI compared to controls and were statistically significant **(Figure 6)**. By contrast, challenge with Td-CF522 failed to recapitulate this effect **(Figure 6)**.

**Figure 6.**
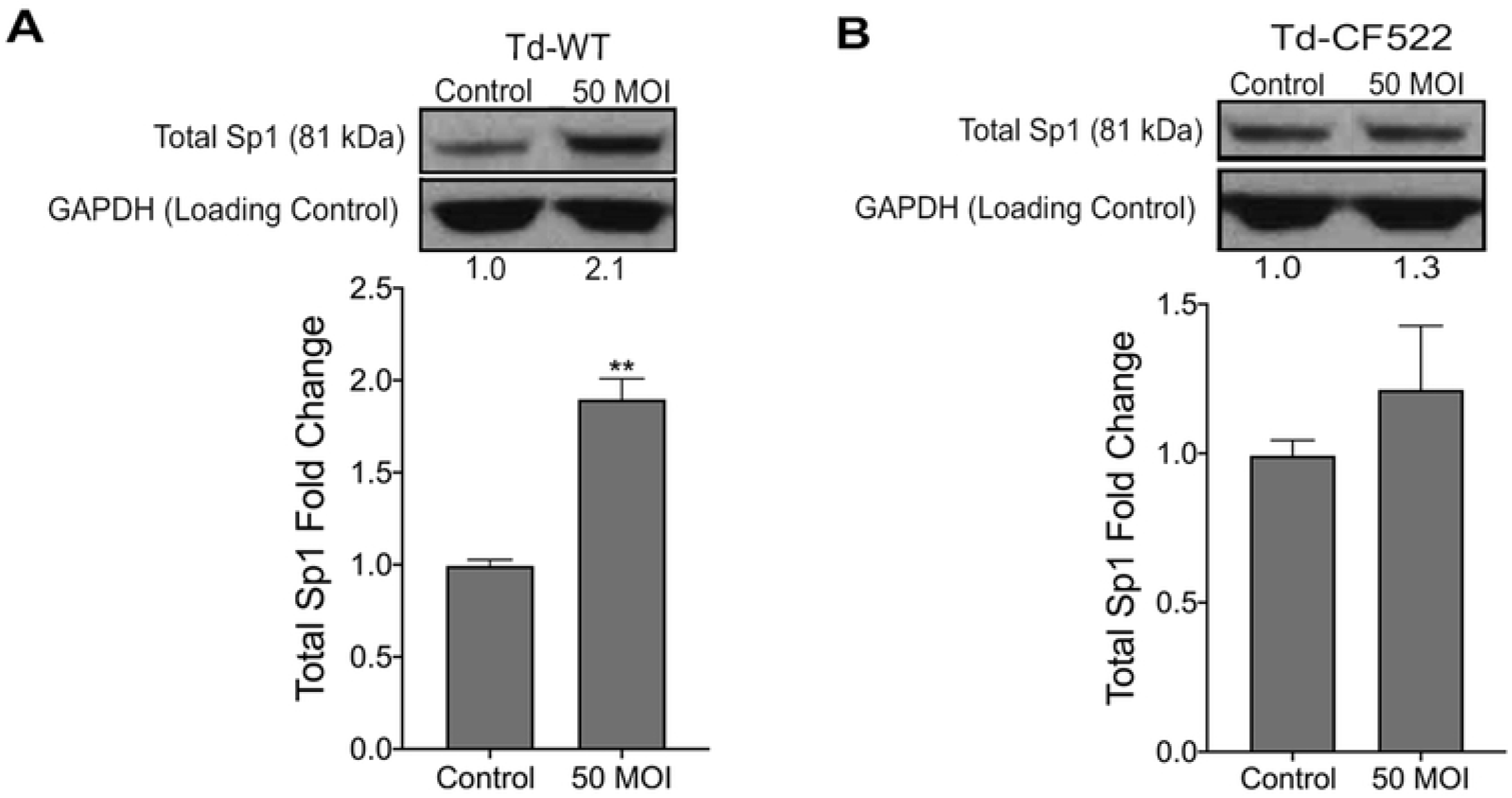
Healthy hPDL cells were challenged with A) Td-WT and B) isogenic Td-CF522 bacteria at an MOI of 50 as previously described. Whole cell lysates were generated and used for Western Blot analysis utilizing total anti-Sp1 antibodies. Total Sp1 protein expression was normalized against GAPDH protein expression as a loading control. Statistical significance was determined using a paired t-test. Bars represent mean ± SD (n=3). ^**^p<.01 versus control.

In the earliest stages of infection, transcription factors are activated post-translationally by TLR signaling, often at the step of nuclear translocation leading to the induction of the primary response genes(76). After finding that *T. denticola* increases total Sp1 protein expression, we investigated whether *T. denticola or dentilisin* are able to induce nuclear translocation of Total Sp1 via TLR2 and MyD88 activation in hPDL cells and the results can be seen on **Figure 7 and 8**. *TLR2, MyD88*-deficient and shRNA control cells were stimulated with purified dentilisin, Td-WT or Td-CF522 as previously described. Protein localization was assessed using primary antibodies against Sp1 and a secondary antibody conjugated to an Alexa 488 fluorophore and subjected to confocal microscopy. shRNA control hPDL cells challenged with Td-CF522 resulted in an increased Sp1 protein signal compared to the control group but remained localized to the cytoplasm **(Figure 7A)**. In contrast, shRNA control hPDL cells stimulated with purified dentilisin and Td-WT bacteria resulted in an increased Sp1 signal and translocation to the nucleus **(Figure 7A)**. hPDL cells deficient in *TLR2* and *MyD88* and challenged with the Td-CF522 resulted in similar results as the shRNA control group **(Figure 7 and 8)**. However, when challenged or stimulated with purified dentilisin and Td-WT bacteria, both *TLR2* and *MyD88* deficient cells inhibited the translocation of Sp1 into the nucleus **(Figure 7 and 8A)**. Thus, after determining both TLR2 and MyD88 play an integral role in the translocation of Sp1 in our system, we sought out to determine if Sp1 is required or sufficient to regulate target MMPs.

**Figure 7.**
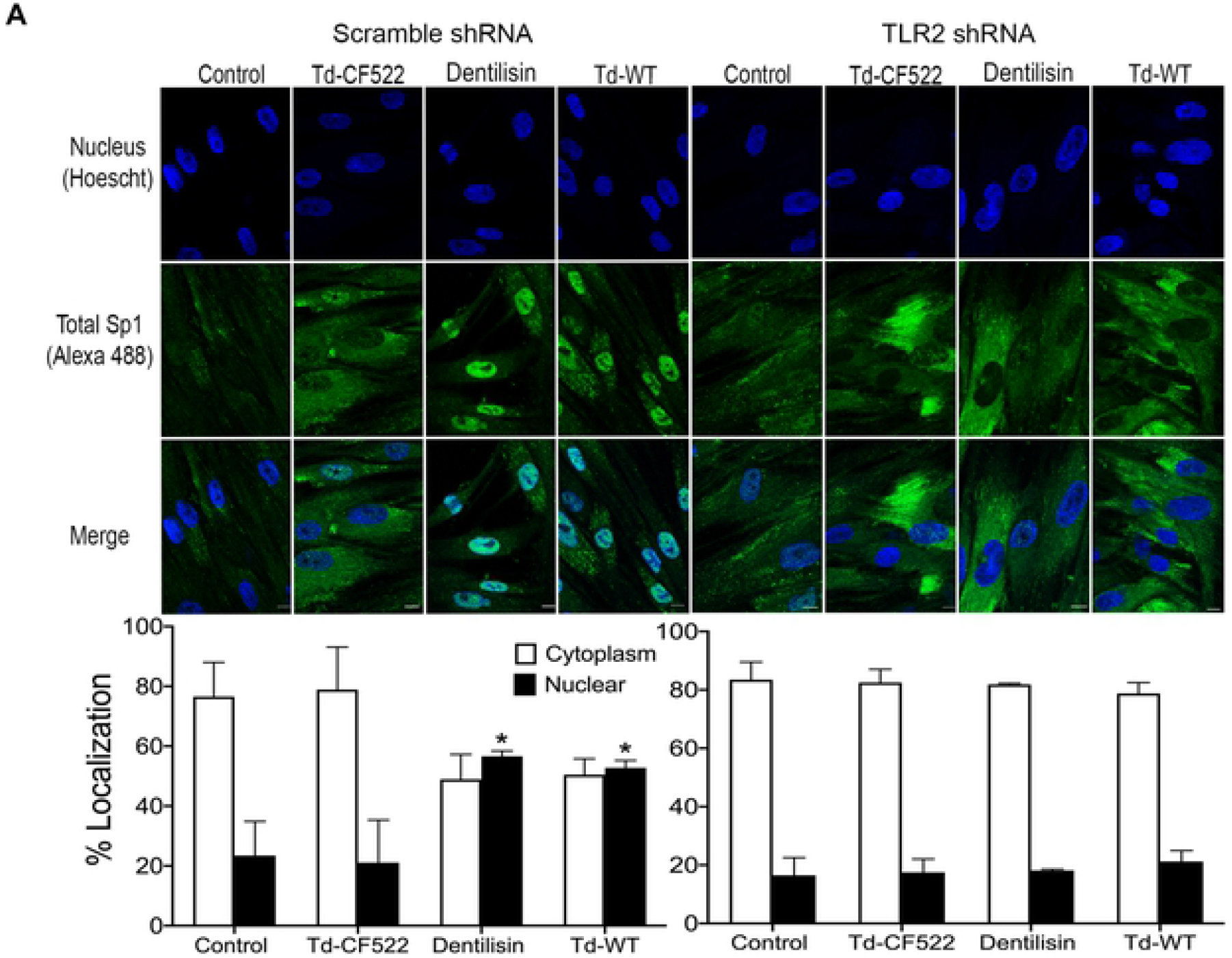
Knockdown of *TLR2*-dependent signaling inhibits translocation of transcription factor Sp1 in both *T. denticola*- and dentilisin-stimulated hPDL cells. A-B) Healthy hPDL cells were challenged with Td-WT and isogenic Td-CF522 bacteria at an MOI of 50 as previously described. Whole cell lysates were generated and used for Western blot analysis utilizing total Sp1-specific antibodies. Total Sp1 protein expression was normalized using GAPDH as a loading control. Statistical significance was determined using a paired t-test. Bars represent mean ± SD (n=3). ^**^*p<*.01 versus control. C) Scramble shRNA control and *TLR2* shRNA hPDL cells were challenged or stimulated with Td-CF522, purified dentilisin or Td-WT at an MOI of 50 and concentration of 1 *µ*g/mL for 2-hours in alpha-MEM media with no supplementation followed by a 22-hour incubation in alpha-MEM media supplemented with 10% FBS, 1% Pen/Strep and 1% Amphotericin B. Cells were stained with Hoescht 33342 (Blue) and total Sp1-specific antibodies (Green), and subjected to confocal microscopy. Scale bar represents 10 *µ*m.

**Figure 8.**
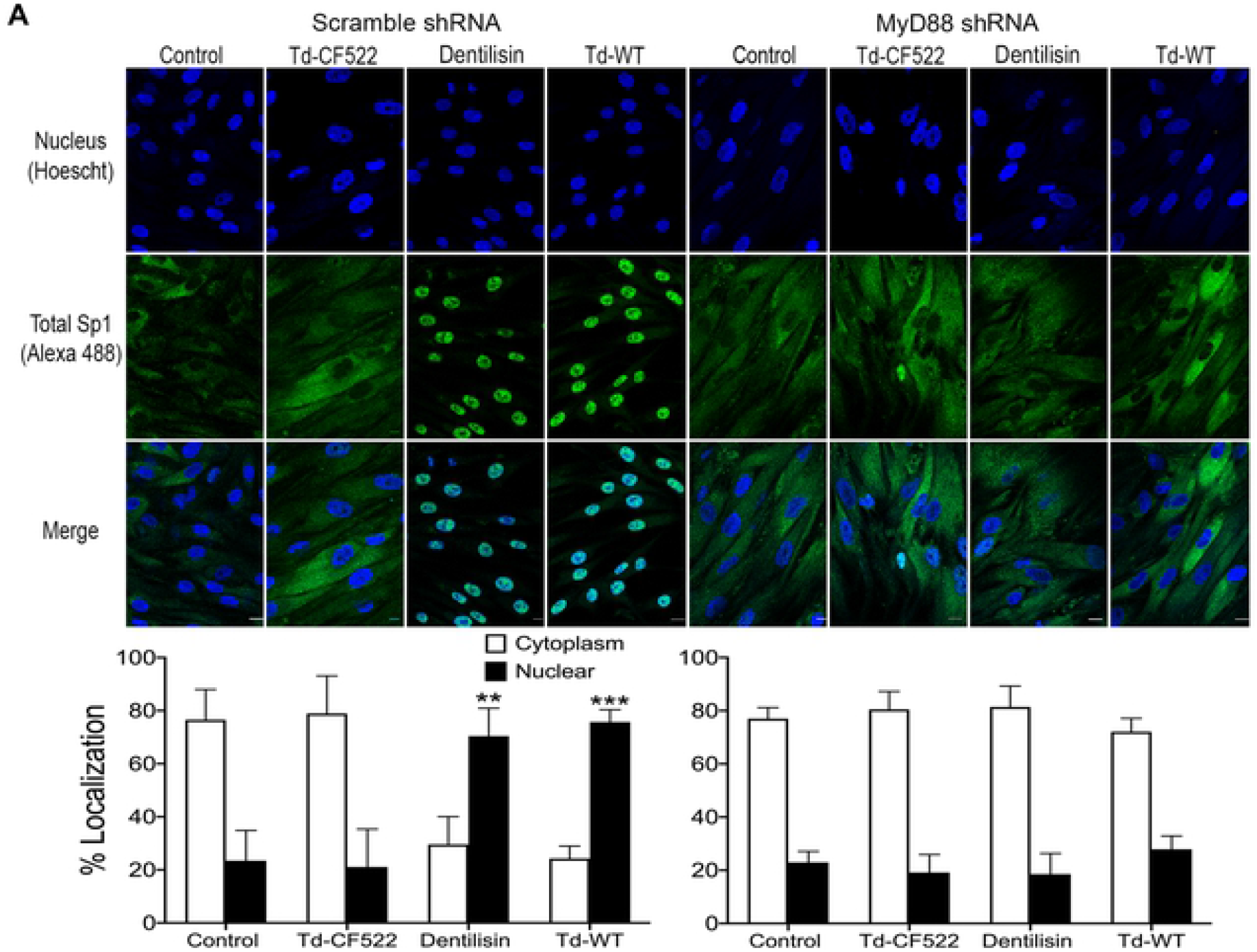
Knockdown of MyD88-dependent signaling inhibits translocation of transcription factor Sp1 in both *T. denticola*- and dentilisin-stimulated hPDL cells. A) Scramble shRNA control and *MyD88* shRNA hPDL cells were challenged or stimulated with Td-CF522, purified dentilisin or Td-WT at an MOI of 50 and concentration of 1 *µ*g/mL for 2-hours in alpha-MEM media with no supplementation followed by a 22-hour incubation in alpha-MEM media supplemented with 10% FBS, 1% Pen/Strep and 1% Amphotericin B. Cells were stained with Hoescht 33342 (Blue) and total Sp1-specific antibodies (Green), and subjected to confocal microscopy. Scale bar represents 10 *µ*m. Scale bar represents 10 *µ*m.

### Knockdown of transcription factor Sp1 inhibits *T. denticola*/dentilisin-stimulated upregulation of MMPs 2, 11, 14, 17 and 28 in hPDL cells

Promoter characterization and luciferase promoter activity assays have identified Sp1 as a regulator of most MMPs and TLRs, through interactions with other transcription factors and cis-regulatory elements such as distal enhancers(77-82). Thus, it is possible that Sp1 may function as a critical regulator in response to *T. denticola* challenge. To answer this question, shRNA vectors against the *Sp1* gene were used to transduce healthy patient-derived hPDL cells using lentiviral particles as previously described. Cells transfected with scrambled shRNA were used as a negative control. Average basal Sp1 mRNA levels were decreased ∼40% across three clones as assessed using RT-qPCR **(Figure 9A)**. These shRNA control and Sp1 deficient hPDL cells were then challenged with purified dentilisin at 1 ug/mL concentration or Td-WT bacteria at a MOI of 50 followed by RNA isolation and RT-qPCR as previously described. Control shRNA hPDL cells stimulated or challenged with purified dentilisin and Td-WT bacteria resulted in the statistically significant upregulation of MMPs 2, 11, 14, 17 and 28 **(Figure 9B-9F)**. By contrast, Sp1 deficient hPDL cells under the same conditions suppressed the upregulation of all MMP targets **(Figure 9B-9F)**.

**Figure 9.**
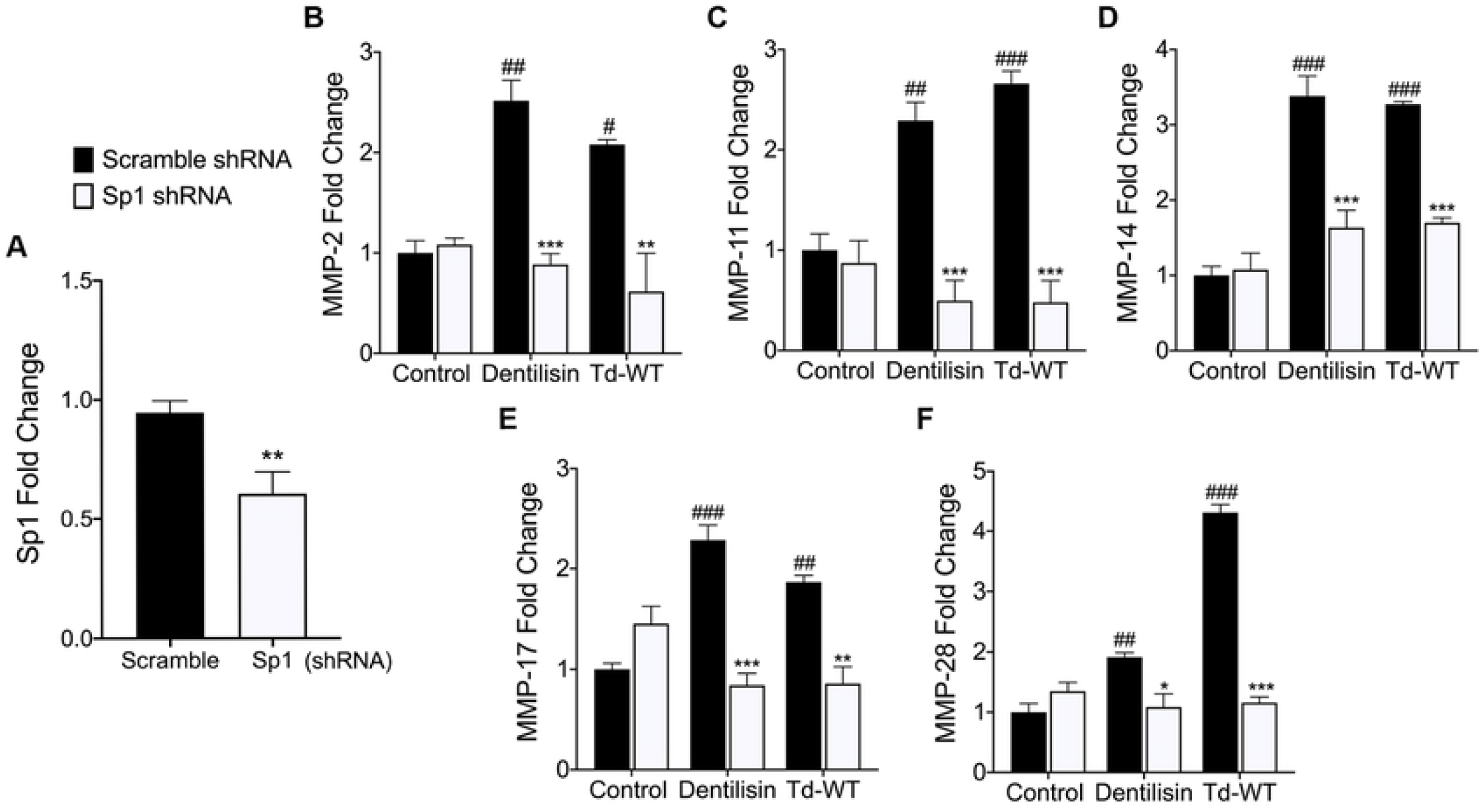
Stable Suppression of *Sp1* inhibits *T. denticola*-stimulated upregulation of MMP *2, 11, 14, 17* and *28* in hPDL cells. **A)** RT-qPCR validation of stable gene suppression using shRNA vectors targeted against *Sp1* in healthy hPDL cells. Cells transduced with scramble shRNA vectors were used as a control. Statistical significance was determined using an unpaired t-test. Bars represent ± SEM of mean values (n=3 clones). ^**^*p*<.01 versus control. **B-F)** RT-qPCR for *MMP-2, MMP-11, MMP-14, MMP-17 and MMP-28* mRNA expression of scramble shRNA control and *Sp1* shRNA hPDL cells challenged or stimulated with purified dentilisin at a concentration of 1 *µ*g/mL or Td-WT at an MOI of 50 for 2-hours in alpha-MEM media supplemented with 10% FBS and no antibiotics followed by a 22-hour incubation in alpha-MEM media supplemented with 10% FBS, 1% Pen/Strep and 1% Amphotericin B. The expression of each gene was normalized to that of *GAPDH*. Statistical significance was determined using a Two-Way ANOVA followed by post-hoc Tukey’s multiple comparisons. Bars represent mean ± SEM (n=3). #*p*<.05 versus scramble control group. ##*p*<.01 versus scramble control group. ###*p<*.001 versus scramble control group. ^*^*p*<.05 versus *Sp1* shRNA equivalent group. ^**^*p<*.01 versus *Sp1* shRNA equivalent group. ^***^*p<*.001 versus *Sp1* shRNA equivalent group.

## DISCUSSION

In this study, we characterized molecular mechanisms of dentilisin-dependent TLR2 activation and MMP expression in hPDL cells, using *T. denticola* (both wildtype and dentilisin mutant) and purified native dentilisin. This robust approach demostrated the biological relevance of dentilisin in modulating key host tissue response pathways that control periodontal tissue homeostasis.

Data from this study extends our earlier reports that short term exposure to dentilisin leads to sustained upregulation of tissue destructive processes at both the transcriptional and translational levels. Specifically, we demonstrated that *T. denticola* acylated dentilisin plays a direct role in activating TLR2/MyD88-dependent pathways resulting in the subsequent upregulation of MMPs in hPDL cells. Additionally, we have identified the transcription factor Sp1 as an important downstream target that is activated as a result of TLR2/MyD88 activation, leading to its translocation into the nucleus and mediating the upregulation of various MMPs **(Figure 10)**.

**Figure 10.**
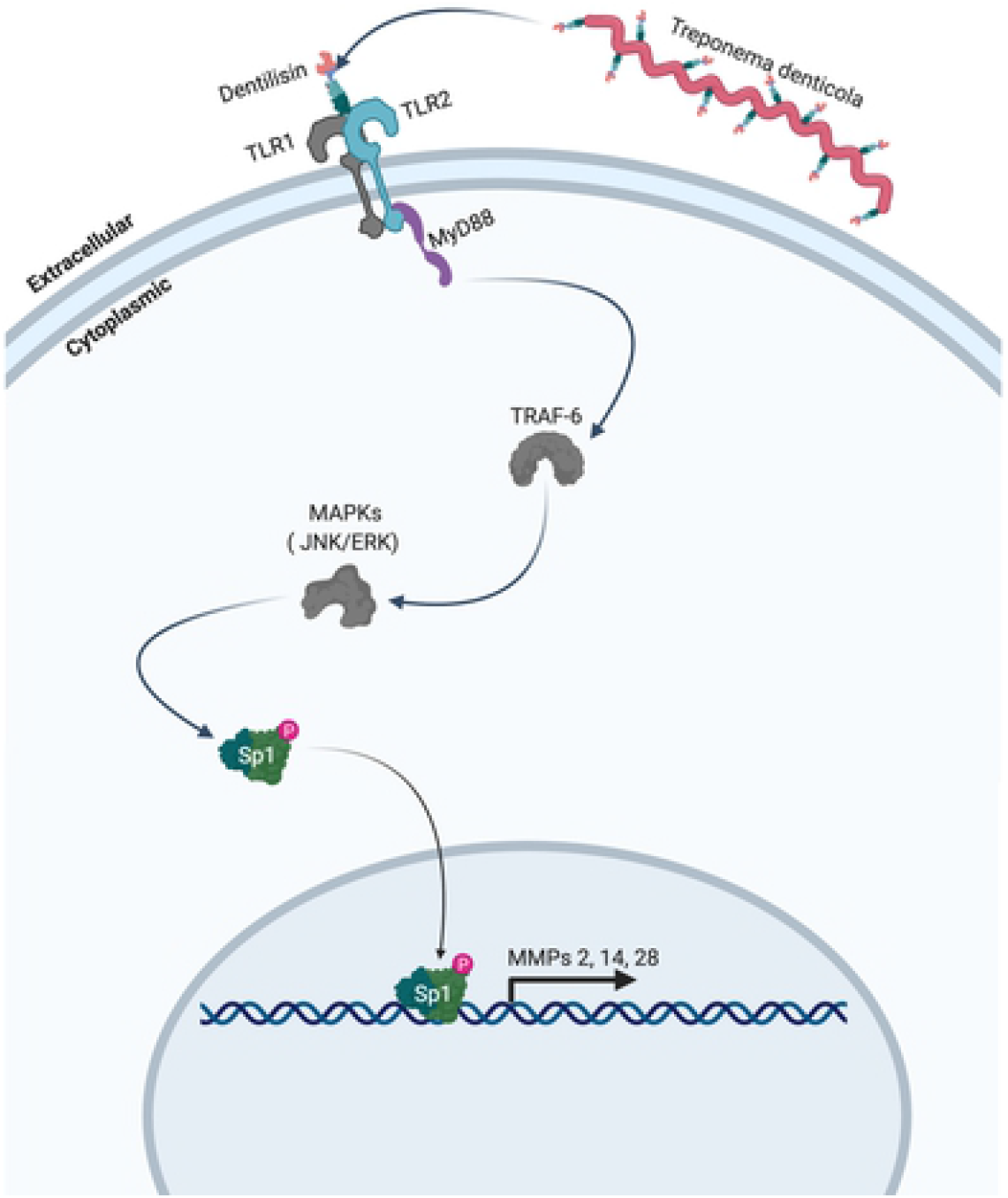
Model of proposed mechanism. *T. denticola* activates host expressed TLR2 receptors via surface-expressed dentilisin. Subsequent downstream activation of MyD88 and translocation of the transcription factor Sp1 lead to the upregulation of MMPs 2, 11, 14, 17 and 28 in human periodontal ligament cells. Grey colored icons represent genes that were identified through RNA-sequencing and analysis but were not directly validated.

The main enzymes responsible for catabolic breakdown of collagen and other ECM proteins are MMPs(83). Further investigations have led to the discovery of a multitude of MMP-dependent phenotypes and novel functions in health and disease(17, 56, 84). As a result, it has become increasingly clear that MMPs act as master regulators of inflammation, through proteolysis of cytokines/chemokines, growth factors, receptors and their binding proteins, proteases, protease inhibitors, as well as intracellular multifunctional proteins(55, 84-87).

Our RNA-Seq analysis revealed specific gene expression patterns that are associated with distinct pathologic features presented clinically(16), reported in *in vivo* studies(88) and previous studies from our lab(41, 44). We recently reported that short term *T. denticola* challenge induced chronic upregulation of MMP-2 and 14 expression *in vitro* as long as 12 days. Our present data expands on this finding demonstrating *T. denticola’s* ability to drive a tissue-destructive gene profile after a brief exposure. While the 5-hour incubation group clustered with the control group, the 24-hour incubation group clustered away from the control group illustrating *T. denticola’s* ability to mediate large-scale transcriptional changes within 24-hours after infection **(Figure 1B)**.

ECM degradation is a key event in the early stages of periodontal disease, generating ECM fragments that can act as danger-associated molecular patterns (DAMPs). ECM fragments that act as DAMPs include fragmented aggrecan, fibronectin, biglycan, decorin and low molecular weight hyaluronic acid(89). Fibronectin fragments have been identified as biomarkers of periodontal disease, with specific fragments (i.e. 40-, 68- and 120 kDa) capable of modulating various cellular processes including apoptosis, proliferation and proteinase expression through integrin signaling(90-94). Although TLRs were originally characterized for their role in innate immunity, recent findings have delineated DAMPs as TLR agonist as well. Hwang *et al*. 2015 demonstrated that 29-kDa fibronectin fragments induced catabolic responses, including MMP expression, by activation of p38 through a MyD88-dependent TLR-2 signaling pathway in articular chondrocyte cultures(92). Other groups have revealed a role of fibronectin in innate immune responses of macrophages through cooperating with TLR2 and TLR4, which lead to the upregulation of pro-inflammatory genes.

We recently reported that *T. denticola* expressed dentilisin indirectly mediates fragmentation of extracellular fibronectin in a host MMP-2 -dependent manner(37). Differential expression analysis of *T. denticola-*challenged hPDL cells revealed fibronectin as the most upregulated ECM constituent in the “ECM Interactions” and “ECM Organization” gene ontology terms with a concomitant enrichment of genes associated with the degradation of various ECM proteins, including MMPs 11, 17 and 28 **(Figure 1C)**. Interestingly, while the upregulation of integrins associated with collagen and laminin interactions were detected, such as α10 and β4, integrins responsible for sensing full-length or fragmented fibronectin did not change. Taken together, these data and reports in literature suggest that *T*.*denticola* dentilisin-induced fibronectin fragmentation may play a direct role in activating and amplifying TLR2-dependent signaling pathways, exacerbating periodontal tissue degradation.

PDLs insert into the cementum through Sharpey’s fibers which are primarily comprised of fibrillar collagen(95, 96). While most collagen is degraded by collagenases such as MMP-1, MMP-8 and MMP-13, MMP-2 is the predominately expressed species in human PDLs and has been shown to diffuse and actively degrade fibrillar collagen(83). MMP proteolytic cascades can also lead to widespread periodontal tissue destruction due to cooperative MMP activation(97). Although MMP-14 has no direct role in degrading collagen, it is a potent activator of pro-enzymatic MMP-2(97). Additionally, both MMP-2 and MMP-14 participate in the active degradation of various cytokine and chemokine substrates expressed by hPDL including IL-1β, IL-6, TNF-α, CXCL12, IL-8 and MCP protein family(55). Lastly, although MMPs are classically viewed as extracellular and localized to the cell surface, they have also been found in every cellular compartment(21). Compelling evidence has revealed MMP-14 moonlighting as a transcription factor independent of its catalytic activity, regulating >100 genes associated with inflammation and bacterial clearance using chromatin immunoprecipitation (ChIP)-sequencing and reporter vector analysis(21, 85). Therefore, it is possible that MMP-14 regulates other MMPs while initiating a forward feed-back loop to regulate itself. Taken together, these data suggest that dysregulation of MMP-2 and MMP-14 play a key role in periodontal disease progression.

While MMPs 2 and 14 have been linked to periodontal disease progression, MMPs 11, 17 and 28 have not(17). Similar to other MMPs, clear roles for MMP-17 (also called MT4-MMP) have been described within the context of cancer(98). However, identifying a specific role in tissue homeostasis and other disease has been rather challenging. The GPI-anchored MMP-17 has minimal catalytic activity against ECM substrates only showing affinity for fibrin/fibrinogen, gelatin and pro-TNF-alpha ligands(98, 99). To date, precise analysis of MMP-17 null mice has revealed a weak phenotype associated with water retention by regulating processes in the hypothalmus(100). Additionally, MMP-17 mediates C-terminal processing of ADAMTS4, one of the aggrecanases that are thought to play a role in arthritis(101). Pro-MMP-11 (also called Stromelysin-3) contains a unique pro-peptide motif for intracellular calcium activation by furin and has also demonstrated rather restricted substrate specificity(102, 103). The most clearly established function of MMP-11 is associated with adipocyte-differentiation and tumor migration through catalysis of IGFBP-(104, 105). MMP-11 ECM substrates identified so far includes the laminin receptor, fibronectin, elastin and the native alpha3 chain of collagen(106); all constituent ECM proteins found to be upregulated in our differential expression analysis. Gene ontology analysis also clustered MMP-11 with various collagen and ECM catabolic processes **(Figure 1C)**, highlighting MMP-11 as a potentially unique response regulator during periodontal disease.

*MMP-28* overexpression *in vitro* lead to increased *MMP-2* and *MMP-14* mRNA levels, constitutive MMP-2 activation, and revealed e-cadherin as a high-affinity substrate for proteasomal degradation (107, 108). Our data aligned with these findings as *MMP-28* was the most upregulated MMP (∼2-fold change) within the proteinaceous ECM gene ontology term and was not expressed in control groups **(Figure 1C)**. Taken together, these data suggest that MMP-28 plays a multifaceted role in exacerbating *MMP-2* and *MMP-14* mRNA upregulation while directly contributing to their constitutive activation during periodontal disease progression. Taken together, the multi-dimensionality of MMPs and their varied localizations highlight their role as integral host factors that mediate immune evasion and inhibit bacterial clearance through a web of complex interactions. Thus, a more thorough characterization of MMP functionality in the context of specific bacterial DAMPs may reveal new therapeutic targets and elucidate novel regulatory mechanisms underpinning periodontal disease.

Although both TLR2 and TLR4 have been implicated in clinical periodontal disease progression, various studies utilizing TLR2 knockout mice and *in vitro* systems suggest that it plays the most direct role is in sensing various components produced by periodontal pathogens underpinning alveolar bone destruction(34, 36, 88, 109). However, the precise roles, whether protective or destructive, played by TLR2 in periodontal infection and inflammation are poorly understood, particularly in the context of *T. denticola* infection. In addition, high-throughput studies and the advent of bioinformatics have demonstrated that a single microbial antigen can be sensed by multiple extracellular and intracellular pattern recognition receptors(110). Thus, multi-TLR cooperation may at least partially explain this differential response. Nevertheless, since previous studies have typically utilized synthetic ligands or LPS, or have limited their analysis to a few outputs, a comprehensive analysis is needed to reveal the intrinsic factor(s) mediating TLR signaling crosstalk and the tissue-specific effects.

Once a bacterial ligand binds, a conformational change is thought to occur that brings the two Toll/interleukin-1 (IL-1) receptor (TIR) domains on the cytosolic face of each receptor into closer proximity, creating a new platform on which to build a signaling complex through various adaptor molecules(111). Current lines of thinking suggest these heterodimerizations of TLR2 are an integral factor differentiating between pro-vs. anti-inflammatory responses(72). These adaptors are MyD88, MyD88-adaptor-like (MAL, also known as TIRAP), TIR-domain-containing adaptor protein inducing IFNβ (TRIF; also known as TICAM1), TRIF-related adaptor molecule (TRAM; also known as TICAM2) and sterile α- and armadillo-motif containing protein (SARM)(112). Thus, availability of intracellular adaptors is another factor that can accommodate distinct signaling patterns as a result of TLR2 activation(72). Because MyD88 is the predominant mediator for almost all TLRs with the exception of TLR3 and TLR4, we sought out to determine its role in regulating MMP expression in PDL tissues. MyD88 deficient hPDL cells stimulated with Td-WT or purified dentilisin resulted in universal suppression of all MMP targets compared to the shRNA control group **(Figure 5)**. Similarly, TLR2 deficiency (in hPDL cells under the same experimental conditions) was sufficient to suppress all MMP targets with the exception of MMP-11 **(Figure 4)**. Taken together, our data suggests that *T. denticola* utilizes a TLR22/MyD88-dependent pathway to regulate MMPs 2, 11, 14, 17 and 28 in PDL tissues.

Several other pathways are able to modulate TLR-signaling, thereby fostering unique and tissue-specific outcomes. Synergistic cooperation between the Notch pathway and acute TLR-induced signals in the activation of canonical and non-canonical Notch target genes has been characterized as an integral pathway governing the regulation of innate and adaptive immunity(113-116). However, mechanisms by which TLRs induce Notch ligand expression and indirectly activate Notch signaling are not well understood. Whether Notch activation is a negative or positive regulator of TLR signaling is unclear and may depend on cell or tissue type(117-119). For activation of classic Notch target genes in macrophages, TLRs induced an IKK- and p38-mediated signal that cooperated with canonical Notch signaling that was dependent on RBP-J serving as an integration point in macrophages(117). Our data are in agreement with these findings, as KEGG pathway analysis revealed Notch signaling (∼2-fold change) as the second most enriched pathway resulting from *T. denticola* challenge in hPDL cells 22-hours post-challenge **(Figure 1D)**. Additionally, differential expression analysis revealed a concomitant upregulation of *MAP2K6* and *MAPK11* (data not shown), both constituents of p38-dependent MAP Kinase signaling implicated with the induction of various pro-inflammatory associated genes resulting from TLR activation(120). While crosstalk between these two pathways in homeostasis and disease is well-studied, less is known about the tissue-specific downstream outcomes and the mechanisms governing them, especially within the context of periodontal disease and MMP regulation.

Spirochetes express a wide range of novel and structurally diverse lipoproteins(121). Many are localized to the outer membrane and determine how these unique microbes interact with their environments, including eliciting pro- and anti-inflammatory responses(122). These membranes are transiently shed into the extracellular space through natural blebbing processes as outer membrane vesicles and exhibit a similar protein profile as the outer membrane while the vesicle lumen mainly contains periplasmic proteins(123). The components of the dentilisin complex (acylated protease PrtP and accessory lipoproteins PrcA and PrcB) have been reported as 3 of the most enriched molecules in *T. denticola* outer membrane vesicles(124). This suggest that their lipid moieties are available to TLR2 recognition complexes. In our study, stimulation with purified dentilisin lead to the upregulation of *MMPs 2, 11, 14, 17* and *28*. By contrast, challenging hPDL cells with isogenic Td-CF522 (which lacks the anchoring PrcB and catalytic PrtP subunits)(125) had no effect **(Figure 4)**. Thus, while we originally hypothesized dentilisin lipid moieties to be the causative drivers of TLR2 activation, our data suggest dentilisin protease activity may be an integral factor responsible for the induction of various MMPs. However, the exact mechanism of action remains to be determined.

At the cellular level, the very basis of inflammation is the deployment of complex molecular mechanisms, including hundreds of genes that are activated within minutes after the primary stimulus(76, 126). Although inflammatory pathways include a core set of genes that are almost invariably activated in most cell types and in response to most inflammatory stimuli, they differ extensively depending on the cell type and tissue in which they are interacting with and the intensity of the trigger(76). Examples include nuclear factor-κB (NF-κB)(76, 126), interferon-regulatory factor (IRF) proteins(127) and Kruppel-Like family transcription factors such as Sp1(128, 129). Although increased Sp1 levels have been implicated as a factor underpinning periodontal disease progression, a thorough characterization of its role in periodontal tissues has not been previously addressed. When healthy hPDL cells were challenged with wild-type *T. denticola*, total Sp1 protein expression increased compared to the control group **(Figure 6A)**. By contrast, hPDL cells challenged with its isogenic mutant, Td-CF522, inhibited this effect **(Figure 6B)**. This finding led us to determine if this increase in protein expression was met with a concomitant increase in Sp1 translocation to the nucleus. To answer this question, TLR2 and MyD88 knockdown patient-derived hPDL cells lines were generated and challenged with wild-type *T. denticola*, Td-CF522 and purified dentilisin followed by staining with primary antibodies against Sp1 and secondary conjugated to an Alexa 488 fluorophore. shRNA control cells stimulated with dentilisin and challenged with Td-WT resulted in Sp1 protein localizing to the nucleus, while TLR2 and MyD88 suppresion under the same conditions inhibited this effect compared to the control group **(Figure 7 and 8)**. Interestingly, a slight increase in cytoplasmic Sp1 was noted in shRNA control and *TLR2* and *MyD88* knockdown groups when challenged with the Td-CF522 isogenic mutant, suggesting that other factors expressed by *T. denticola* may also be influencing the regulation of Sp1 expression, whereas dentilisin preferentially regulates its post-translational activation.

Many MMPs, including MMPs 2, 11, 14, 17 and 28, have multiple GC boxes in their proximal promoters which bind Sp1 and potentially other GC-binding proteins(81). MMPs, such as MMP-2 and MMP-14 without other obvious promoter features are generally expressed in a more constitutive fashion(81). For example, the human MMP-14 promoter is crucial in maintaining expression of this gene, since introducing mutations reduces promoter activity by approximately 90%(130). Additionally, the myriad of post-translational modifications that Sp1 is subject to can modulate protein levels, DNA-binding activity, transactivation potential, and recruitment of chromatin remodelers(80). Swingler et al. 2010 reported that Sp1 transcription factors are acetylated in response to HDAC inhibitors, bind to the conserved GT-box of the MMP28 promoter, potentially working in conjunction with p300/CREB to upregulate mRNA expression(78). Our data also suggests that Sp1 plays an integral role in the upregulation of MMPs in the PDL, particularly under *T. denticola* challenge conditions. While shRNAcontrol hPDL cells stimulated with Td-WT and dentilisin resulted in the upregulation of MMPs 2, 11, 14, 17 and 28, Sp1 deficient hPDL cells under the same conditions suppressed this effect **(Figure 9)**. Sp1 has also demonstrated the capacity to regulate cis-regulatory elements that are highly susceptible to cell stimulation(131). This may explain how constitutively expressed MMPs, such as MMP-2 and MMP-14, are still subject to upregulation upon *T. denticola* stimulation. Taken together, these data suggest that the transcription factor Sp1 is an important downstream early response gene which mediates *T. denticola*-induced upregulation of various MMPs in hPDL cells.

This study has potential limitations. Because our samples are generated from human patient tissues, genetic variation may lead to heavily variegated responses. Due to a low sample size, our RNA-Seq in particular may be subject to read count bias and confounding that may have influenced our results. The Pearson’s correlation reflects the linear relationship between two variables accounting for differences in their mean and SD. The more similar the expression profiles for all transcripts are between two samples, the higher the correlation coefficient will be. As shown in **Supplemental Figure 3**, our samples clustered well, suggesting that dispersion effects are low across the 3 biological replicates. In addition, comprehensive analyses have shown that data sets with very low dispersions, a power of 0.8 is easily reached with very low sample size and sequencing depth(132). While our results provide a strong support for mechanisms of action underpinning periodontal disease, other systemic factors could potentially modulate true phenotypic effects. As such, simulating similar experimental conditions in an animal model is necessary to further investigate.

Most studies characterizing *T. denticola* activation of the innate immune system utilize pro-inflammatory cytokine/chemokines as readouts of activation. Studies that focus on the effects of unchecked TLR signaling and its contributions to the dysregulation of tissue destructive genes in periodontal ligament connective tissues are lacking. Considering the limitations of the study, our data in combination with findings in literature suggest that *T. denticola* dentilisin activates TLR2/MyD88-dependent activation leading to subsequent activation of the transcription factor Sp1 to upregulate MMPs 2, 11, 14, 17 and 28 in human PDL tissues. To the best of our knowledge, this is the first study associating dentilisin with TLR2 activation using an *in vivo* isogenic mutant bacteria in conjunction with purified product from its parent strain. Additionally, we also identified potentially tissue-specific inducible MMPs that that may play novel roles in modulating host immune responses that have yet to be characterized in the context of periodontal disease. Therapies targeted to the inhibition or suppression of MMPs could serve as a useful adjunct to periodontal therapy, such as scaling and root planning(16, 84). Gaining a better understanding of the transcriptional and epigenetic components underpinning their regulation in the context of periodontal disease may be useful to generate more efficacious alternative therapeutics in the future.

## Materials and Methods

### Human Subjects Research

Approval to conduct human subjects’ research, including protocols for the collection and use of human teeth and PDL tissue was obtained from the University of California San Francisco Institutional Review Board (#16-20204; reference #227030). Consent was not obtained because data were analyzed anonymously.

### Human PDL Cell Cultures

As described previously, the primary culture of PDL cells was conducted via the direct cell outgrowth method by isolating cells from the PDL tissue around the middle third of extracted healthy human teeth(133, 134). Cells were maintained in Minimum Eagle Medium-α (MEM-α) (Gibco, USA) augmented with 10% fetal bovine serum (FBS) (Gibco, USA), 1% penicillin/streptomycin (P/S), and 1% amphotericin B (Gibco, USA) in a Steri Cycle 370 incubator (Thermo-Fisher Scientific, USA) with a humid atmosphere containing 95% air and 5% CO_2_ at 37 °C. Cell outgrowths were passaged before reaching confluency. Only cells passage three to seven were used for experimentation. Cells were validated by morphological assessment and gene expression of confident biomarkers such as *Periostin*.

### Anaerobic Bacterial Cultures

*T. denticola* ATCC 35405 (Td-WT) and its isogenic mutant, Td CF522 isogenic dentiilisin-deficient mutant Td-CF522 (62).” For your purposes, this is the relevant information on this strain(125) were grown in tanks filled with an anaerobic gas mixture at 37 °C in Oral Treponeme Enrichment Broth (OTEB) (Anaerobe Systems, USA) as previously described(135). Samples were sent to Novogene (USA/China) for paired-end 16S sequencing on a NovaSeq 6000 platform (Illumina, USA) and analysis (Data not shown). *Veilonella parvula* ATCC 10790 was grown under the same anaerobic conditions as *T. denticola* in brain heart infusion media supplemented with 0.075% sodium thioglycolate, 0.1% Tween 80 and 1% of 85% lactic acid and pH adjusted to 6.5-6.6 using 1 M NaOH. Colony morphology on BHI plates were used to confirm purity of cultures.

### Purification of *T. denticola* dentilisin

The dentilisin protease complex was purified from phase-partitioned *T. denticola ATCC 35405* Triton X-114 extracts as described previously(37). Aliquots of the purified dentilisin were stored at −80°C. Activity of purified dentilisin samples was monitored periodically, using the chromogenic substrate succinyl-L-alanyl-L-alanyl-L-prolyl-L-phenylalanine-*p*-nitroanilide (SAAPFNA) as described previously(48).

### Bacterial Challenging of hPDL Cells

∼9×10^5^ healthy hPDL cells were seeded on 6 cm plates. On the next day, these cells were challenged with either *T. denticola*, Td-CF522 or *V. parvula*, as previously described by Ateia et al.(44). Briefly, the bacteria were centrifuged at 4000 RPM for 15 minutes and the supernatant was removed. The pelleted bacterial cells were then resuspended in antibiotic-free MEM-α (without phenol red) at an optical density (OD) of 0.1 at 600 nm using a Spectramax M2 microplate spectrophotometer (Molecular Devices, USA). It has been previously established that an OD of 0.1 at 600nm is equivalent to 2.4×10^8^ CFU/ml for *T. denticola(135)* and 2×10^8^ CFU/mL for *V. parvula* strains*(136)*. Next, hPDL cells were challenged with the bacteria at 50 multiplicity of infection (MOI), whereas the control group was challenged with antibiotic-free MEM-α. Next, the cells were incubated for 2h at 37 °C and 5% CO_2_, washed twice with Phosphate Buffer Saline (PBS) (Gibco, USA) and incubated again at 37 °C and 5% CO_2_ overnight in antibiotic-free MEM-α. On the next day, cells were either harvested for RNA isolation, generation of cell lysates and stored at −80 °C for further investigations.

### RNA-Seq

RNA was extracted from ∼9.5×10^5^ hPDL cells challenged with wild-type *T. denticola* (as described above using an RNeasy Mini Kit and following the manufacturers protocol (Qiagen, Germany). The quality of the extracted RNA was assessed using a Nanodrop UV-Vis Spectrophotometer (Thermo-Fisher Scientific, USA) and by calculating the percentage of RNA fragments with size > 200 bp (DV200) using an Agilent 2100 Bioanalyzer. The RNA was used for first and second strand synthesis, polyA tail bead capture, and sequencing adapter ligation using a TruSeq RNA Library Prep Kit v2 (Illumina, USA). Libraries were sent to Novogene Genomic Services (Davis, USA) for paired-end sequencing on a HiSeq 4000 instrument (150 bp paired-end reads) (Illumina, USA) and analysis. The mean gene expression across all replicates were used for data visualization. Analysis was conducted using the opensource statistical software called R(137) and figures were produced using the Grammar of Graphics Plot 2 (GGplot2)(138) software package.

### Quantitative Reverse Transcriptase Polymerase Chain Reaction (qRT-PCR)

Total hPDL RNA was extracted and purified using the RNeasy mini kit (Qiagen, Germany) according to the manufacturer’s instructions and quantified using a NanoDrop UV-Vis Spectrophotometer (Thermo Scientific, USA). Reverse transcription of the RNA into cDNA was then performed using the SuperScript III cDNA synthesis kit (Cat #11754050, Invitrogen, USA). Samples were ran on a Bio-Rad MyCycler Thermal Cycler according to manufacturers protocol. All cDNA samples were diluted to 5 ng/mL working concentrations and stored at −20°C. cDNA samples were probed for target gene expression via RT-qPCR using PowerUp SYBR Green Master Mix (Applied Biosystems, USA) and primer sequences (Table 1) on a QuantStudio 3 platform (Applied Biosystems, USA). The relative expression levels of target genes were plotted as fold change compared with untreated or negative controls. The 2^-ΔΔ^CT method was used to determine relative change in target gene expression, and gene expression was normalized against GAPDH expression.

**Table 1.**
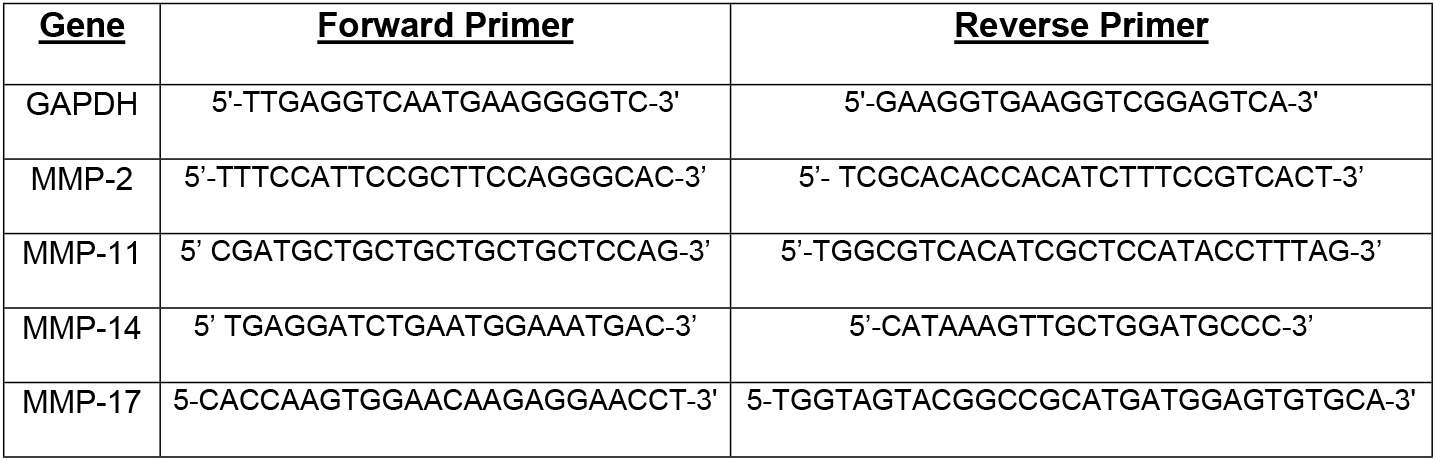
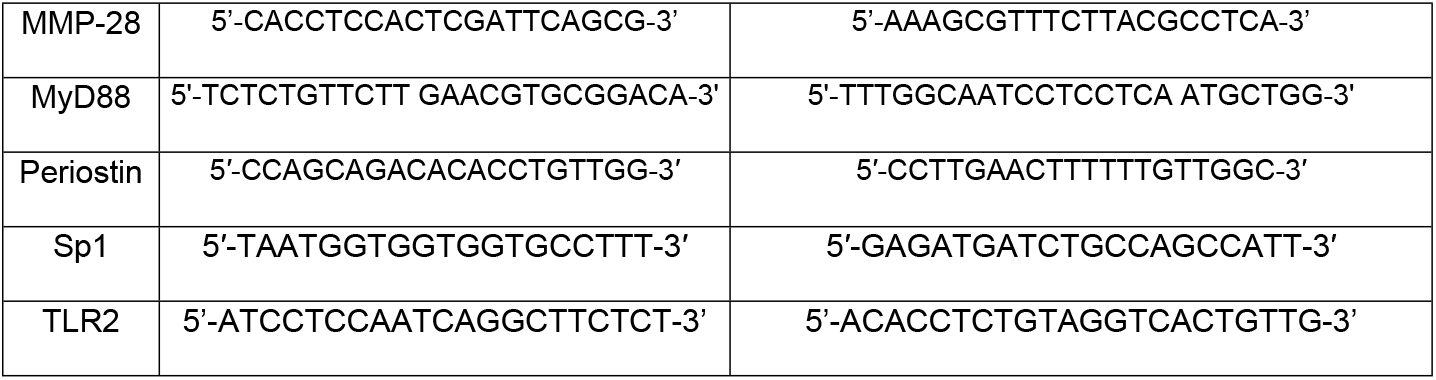
RT-qPCR forward and revers primer pairs

### Stimulation of hPDL Cells Using Purified Dentilisin

A BCA protein assay kit (Thermo Scientific, Rockford, IL, USA) was used to determine purified dentilisin sample concentrations as described above. Assessment of enzymatic activity was determined using gelatin zymography as described below. ∼9.5×10^5^ cells were seeded on 6 cm plates. Purified dentilisin was added to MEM-α media with 10% FBS and 1% P/S at a final concentration of 1 *µ*g/mL and added to healthy hPDL cell cultures for 2-hours. These cells were incubated overnight in MEM-α media free of FBS and supplemented with 1% P/S followed by collection of Supernatant (conditioned media) and total RNA extraction (Cat# 74134, Qiagen, Germany). The resulting samples were stored at −80°C until analyzed further in downstream applications.

### Gelatin Zymography

Supernatant collected from dentilisin stimulated hPDL cells were quantified and normalized to an albumin standard using the Pierce™ BCA protein assay kit (Cat# 23225, Thermo Scientific, USA) according to the manufacturers protocol. Equivalent protein concentrations from samples were mixed with nonreducing sample buffer (0.25 M tris-base, 40% glycerol, 0.8% sodium dodecyl sulfate (SDS), and 0.05% bromophenol blue stain in distilled deionized water/ddH2O at pH 6.8) and loaded into 8% polyacrylamide gels that were co-polymerized with 0.4% SDS and 0.2% gelatin. Samples were electrophoretically resolved on gelatin-containing gels at 125 V for 110 min at 4 °C using a Power Pack supply (Bio-Rad, USA) Next, gels were washed in a series of buffers to facilitate re-constitution of endogenous protein and stained with Coomassie Blue as previously described(44). Briefly, Gels were then washed twice for 15 min under continuous agitation using renaturation/ washing buffer (2.5% v/v Triton-X100 and 0.05 M Tris-base in ddH2O at pH 7.5) to eliminate SDS and promote the renaturation of MMP enzymes. Subsequently, gels were incubated in developing/incubation buffer (0.05MTris-base, 0.15Msodium chloride, 0.01Mcalcium chloride, and 0.02% sodium azide in ddH2O at pH 7.5) for 30 min under agitation, and then the buffer was replaced and gels incubated for 16–20 hr at 37 °C. After that, gels were stained using filtered Coomassie Brilliant blue stain for 1 or 2 hr under agitation. Destaining of the gels was performed using amethanol/acetic acid destaining buffer (40% methanol and 10% acetic acid in ddH2O) until the bands on the gel appeared clear. Zymograms were scanned using an Hp Officejet 6700, and the densitometry of gelatinolytic activity represented by the clear bands was analyzed using Fiji software(139). Brightness and contrast levels of zymogram images were slightly adjusted for publication only.

### Lentiviral shRNA Knockdown

∼6×10^4^ hPDL cells were seeded on 96-well plates. 48 hours later, these cells were infected with either TLR2, MyD88, or Sp1 targeted short hairpin RNA (shRNA) constructs via lentiviral particles according to the kits instructions (Cat# sc-40256-V, Cat# sc-44313-V and Cat# sc-29487-V, Santa Cruz Biotechnology, USA,). Control samples were transduced using nonspecific scrambled shRNA control constructs (Cat# sc-108060, Santa Cruz Biotechnology, USA). hPDL cells were concomitantly permeabilized using Polybrene reagent at a final concentration of 5 µg/mL (EMD Millipore, USA). Approximately 10 hours after infection, cells were washed and incubated overnight in complete MEM-α media supplemented with 10% FBS, 1% PenStrep and 1% Amphotericin B. A pol III promoter drives expression of a silencing cassette containing a puromycin resistance gene used for selection(140). As a result, a short hairpin RNA complex is exported into the cytoplasm, processed by Dicer and assembled into the RISC complex(140). Degradation of targeted mRNA transcripts is mediated by unwinding the siRNA duplex leading to a stable reduction in gene expression. Selection of clones with successfully integrated knockdown constructs was accomplished by treating infected hPDL cells with puromycin (Calbiochem, USA) at a final concentration of 5 µg/mL for 24-hours in supplemented MEM-α media. This media was then exchanged for supplemented MEM-α media containing 1 µg/mL puromycin until single cell colonies were isolated. Finally, cell colonies were expanded and validated via RT-qPCR and/or Western Blot.

### Western Blot Analyses

Cell lysates generated from shRNA lentiviral infection were normalized to an albumin standard as described above. These samples were then subjected to SDS-PAGE 4–12% polyacrylamide gels (Invitrogen, USA) and transferred to polyvinylidene difluoride (PVDF) membranes (EMD Millipore, USA) using a Turbo Blot transfer system and PowerPac Basic power supply (Bio-Rad, USA). The membranes were exposed to a rabbit polyclonal anti-Sp1 (Cat# PA5-29165, Invitrogen, Carlsbad, CA, USA) primary antibody diluted 1:500 in Tris-buffered saline, 0.1% Tween 20 (TBST) solution overnight at 4°C followed by a horseradish peroxidase-conjugated mouse anti-rabbit IgG secondary antibody (Cat# sc-235, Santa Cruz, Dallas, TX, USA) diluted 1:3000 in TBST for 1 hour at room temperature with agitation. Western blots using anti-GAPDH antibodies (Cat# sc-32233, Santa Cruz Biotechnology, USA) were used to confirm equal protein loading. Blots were developed using the Super Signal West Pico kit (ThermoFischer Scientific, USA), scanned for digitization using an Officejet 6700 scanner (Hewlett-Packard, USA) and analyzed using Fiji software(139).

### Confocal Microscopy

Approximately 8×10^4^ hPDL cells were seeded into 4-well glass bottom wells. Cells were challenged with purified dentilisin, wild-type *T. denticola* or isogenic Td-CF522 mutant bacteria as described above. Cells were washed with PBS before being fixed using 4% paraformaldehyde for 10 minutes at room temperature. Next, cells were incubated for 20 minutes with 0.1 M glycine in PBS, washed with PBS and permeabilized with 0.2% Triton X-100 in PBS for 2 minutes at room temperature. Cells were incubated in a 10% serum/PBS blocking buffer for 5 minutes at room temperature followed by a 60-minute incubation in a rabbit sourced polyclonal anti-Sp1primary antibody diluted 1:500 in 10% serum/PBS solution. Next, cells were washed 3 times with PBS followed by incubation in 10% serum/PBS blocking buffer for 1 minute at room temperature followed by incubation in anti-rabbit, mouse secondary antibody conjugated to Alexa 488 fluorophore for 30 minutes at 37°C (1:3000 dilution) and Hoescht 33342 nuclear staining (1:2000 Dilution) in 10% serum/PBS solution. Finally, cells were washed 3 times with PBS, imaged using an SP8 confocal microscope (Leica, Germany) and analyzed using Fiji(139).

### Statistical Analysis

Statistical analysis of differentially expressed genes in RNA-Seq data was assessed using a Kolmogorov-Smirnov test followed by Benjamini-Hochberg correction (*p*<0.05) using R open source software(137). All other data was analyzed using GraphPad Prism Version 8 software (San Diego, USA). Results were evaluated by a one-way ANOVA when comparing more than two groups with a single independent variable while a two-way ANOVA was used to compare more than two groups with 2 independent variables. Western blot and gelatin zymography data was analyzed using a paired or unpaired t-test, respectively.

## DATA AVAILABILITY

The data underlying the results presented in the study are available from https://figshare.com/s/146306b9ef4a7468ed8f

## ACKNOWLEDGMENTS

We would like to thank Charles Le for his technical assistance recovering and maintaining *Veilonella parvula* bacterial stocks.

## FUNDING

This work was supported by funding from National Institutes of Health (R01 DE025225 to YLK and JCF). https://www.nih.gov The funders had no role in study design, data collection and analysis, decision to publish, or preparation of the manuscript.

## COMPETING INTERESTS

*The authors have declared that no competing interests exist*.

## FIGURE LEGENDS

**Supplemental Figure 1.**
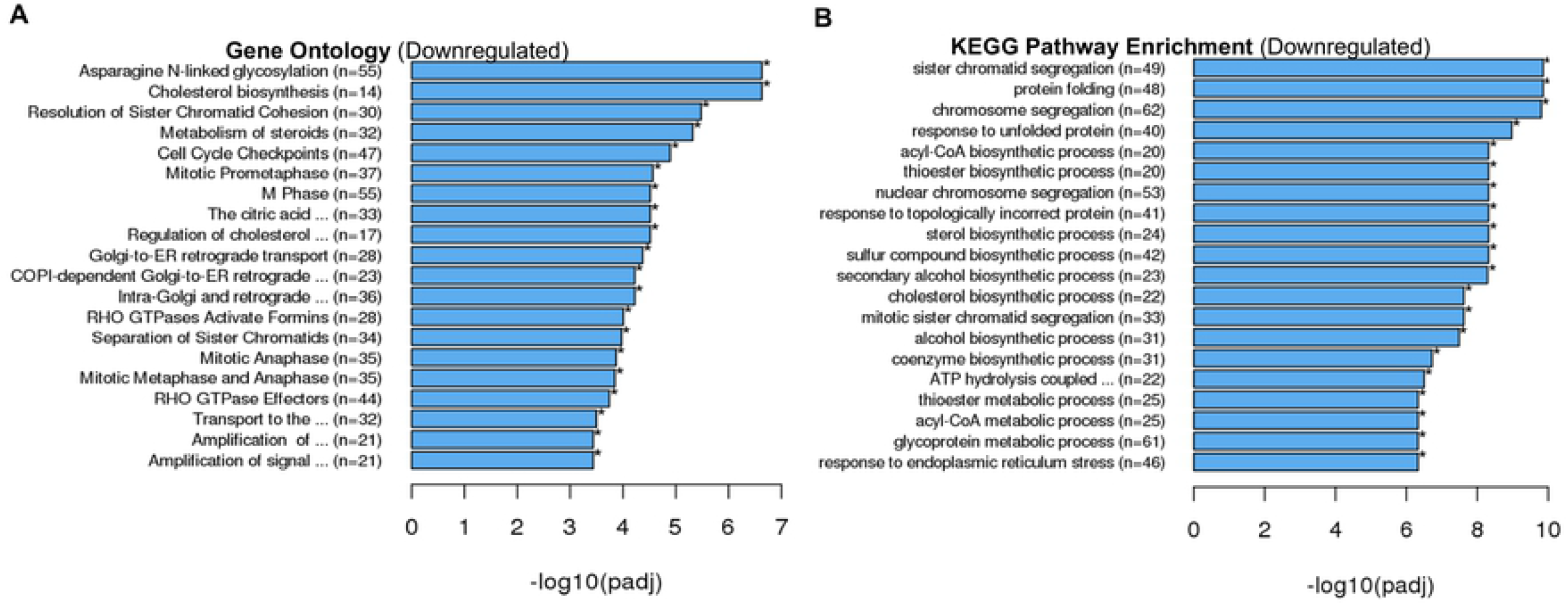
A) Top 20 downregulated Gene Ontology terms of hPDL cells challenged for 2-hours followed by a 22-hour incubation using the Reactome nomenclature. Statistical significance was assessed using a Kolmogorov-Smirnov test followed by Benjamini-Hochberg correction (p<0.05). B) Top 20 downregulated signaling pathways of hPDL cells challenged for 2-hours followed by a 22-hour incubation using the Kyoto Encyclopedia of Genes and Genomes (KEGG) database. Statistical significance was assessed using a Kolmogorov-Smirnov test followed Benjamini-Hochberg correction (p<0.05).

**Supplemental Figure 2.**
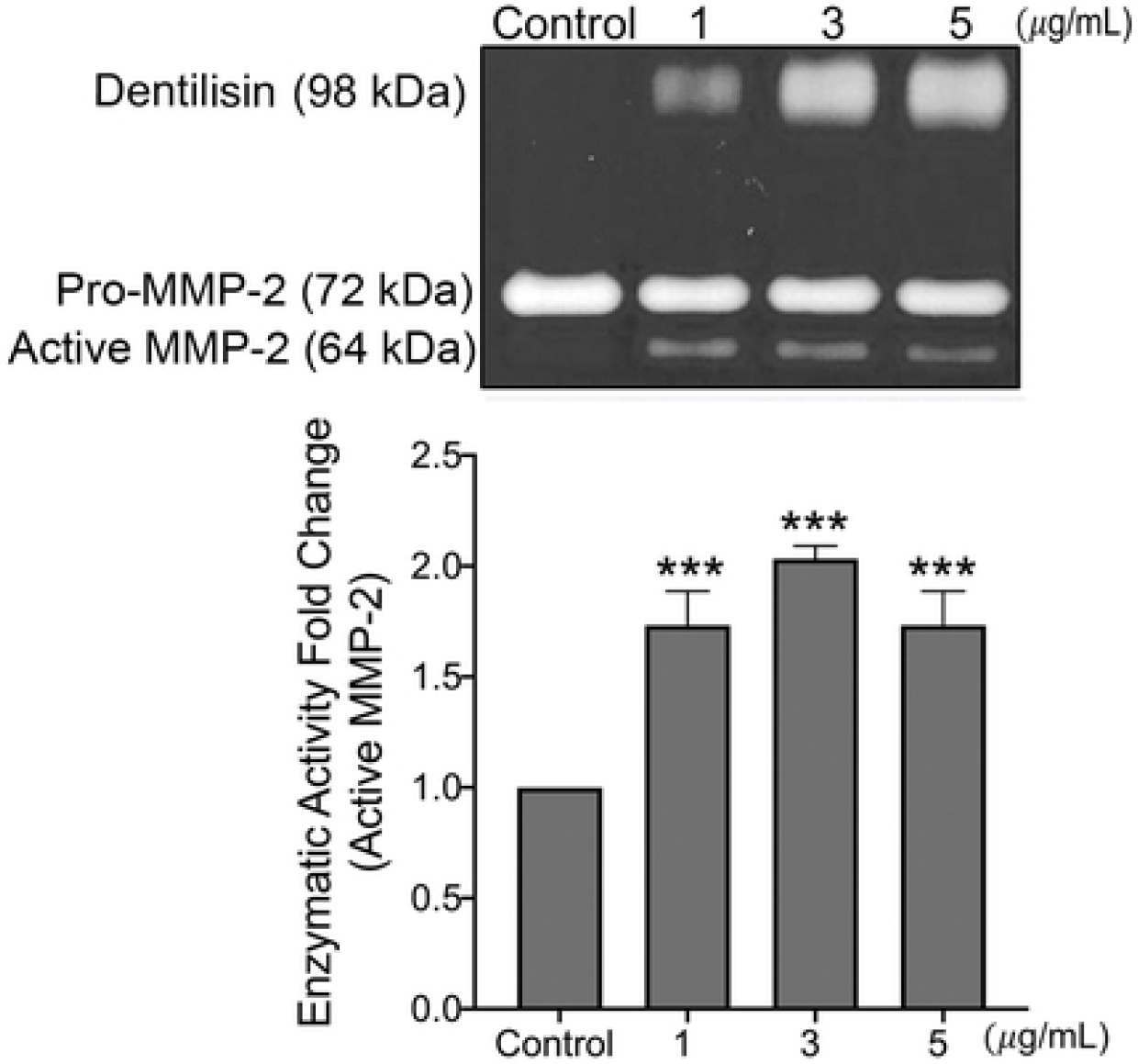
Assessment of purified dentilisin enzymatic activity after stimulation with purified dentilisin. Healthy hPDL cells were challenged with purified dentilisin at increasing concentrations (1, 3 and 5 *µ*g/mL) for 2-hours followed by a 22-hour incubation in MEM-α media free of FBS and supplemented with 1% P/S. Conditioned media from these cells were used to assess the enzymatic activity of Active MMP-2 (64-kDa) and Dentilisin (98 kDa) using gelatin zymography followed by densitometry analysis using Fiji. Statistical significance was determined using a One-Way ANOVA. Bars represent ± SD of mean values (n=3). ^***^p<.001 versus control.

**Supplemental Figure 3.**
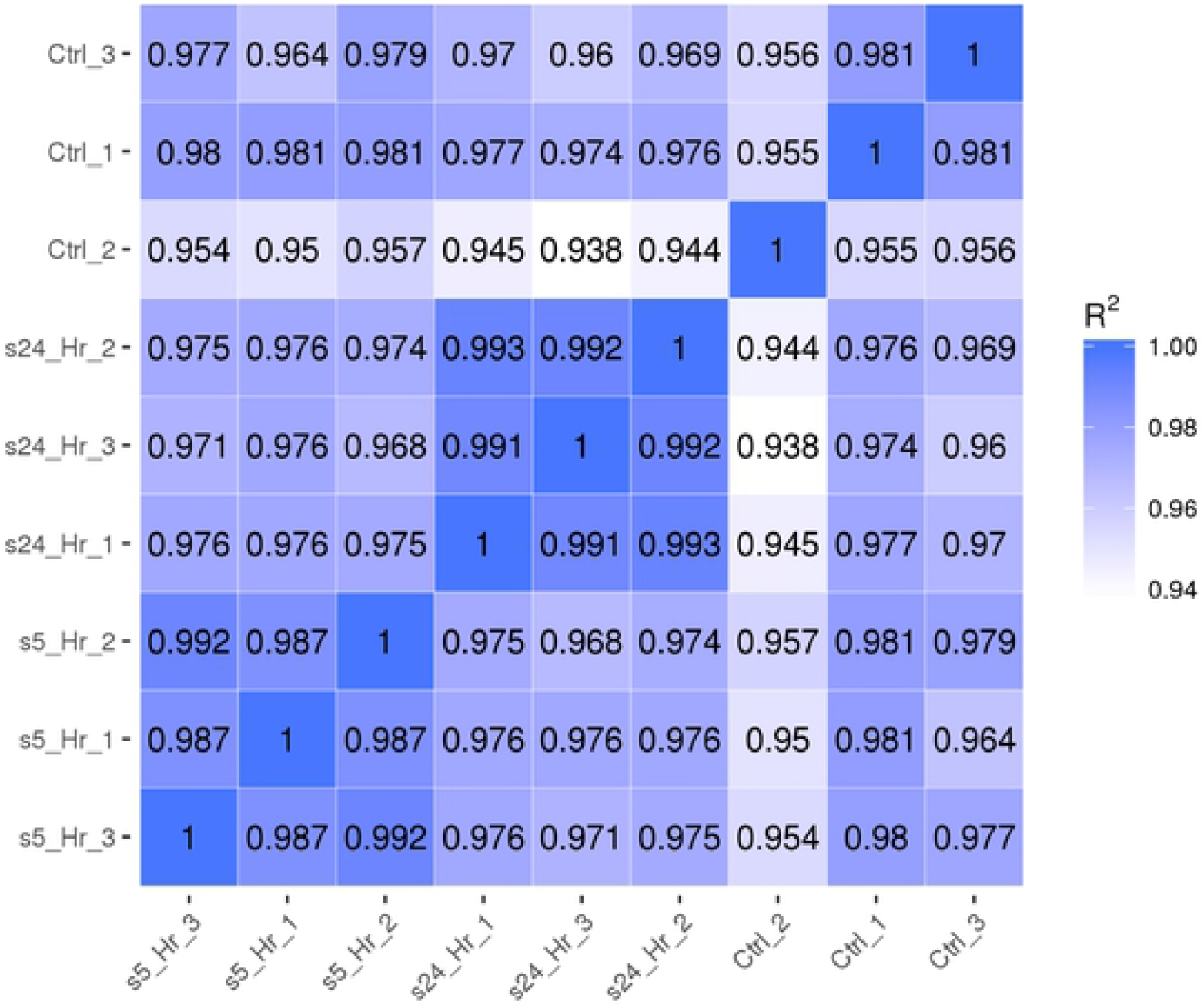
Assessing inter- and intragroup gene expression variability across sample replicates. The figure displays a Pearson’s plot visualizing the correlation between samples. Scale bar represents the range of the correlation coefficients (*R*) displayed.

